# Marginated neutrophils in the lungs effectively compete for nanoparticles targeted to the endothelium, serving as a part of the reticuloendothelial system

**DOI:** 10.1101/2024.06.07.597904

**Authors:** Marco E. Zamora, Eno-Obong Essien, Kartik Bhamidipati, Aditi Murthy, Jing Liu, Hyunjun Kim, Manthan N. Patel, Jia Nong, Zhicheng Wang, Carolann Espy, Fatima N. Chaudhry, Laura Ferguson, Sachchidanand Tiwari, Elizabeth Hood, Oscar A. Marcos-Contreras, Serena Omo-Lamai, Tea Shuvaeva, Evguenia Arguiri, Jichuan Wu, Lubica Rauova, Mortimer Poncz, Maria C. Basil, Edward Cantu, Joseph D. Planer, Kara Spiller, Jarod Zepp, Vladimir Muzykantov, Jacob Myerson, Jacob S. Brenner

## Abstract

Nanomedicine has long pursued the goal of targeted delivery to specific organs and cell types but has not achieved this goal with the vast majority of targets. One rare example of success in this pursuit has been the 25+ years of studies targeting the lung endothelium using nanoparticles conjugated to antibodies against endothelial surface molecules. However, here we show that such “endothelial-targeted” nanocarriers also effectively target the lungs’ numerous marginated neutrophils, which reside in the pulmonary capillaries and patrol for pathogens. We show that marginated neutrophils’ uptake of many of these “endothelial-targeted” nanocarriers is on par with endothelial uptake. This generalizes across diverse nanomaterials and targeting moieties and was even found with physicochemical lung tropism (i.e., without targeting moieties). Further, we observed this in *ex vivo* human lungs and *in vivo* healthy mice, with an increase in marginated neutrophil uptake of nanoparticles caused by local or distant inflammation. These findings have implications for nanomedicine development for lung diseases. These data also suggest that marginated neutrophils, especially in the lungs, should be considered a major part of the reticuloendothelial system (RES), with a special role in clearing nanoparticles that adhere to the lumenal surfaces of blood vessels.

**Graphical Abstract:** 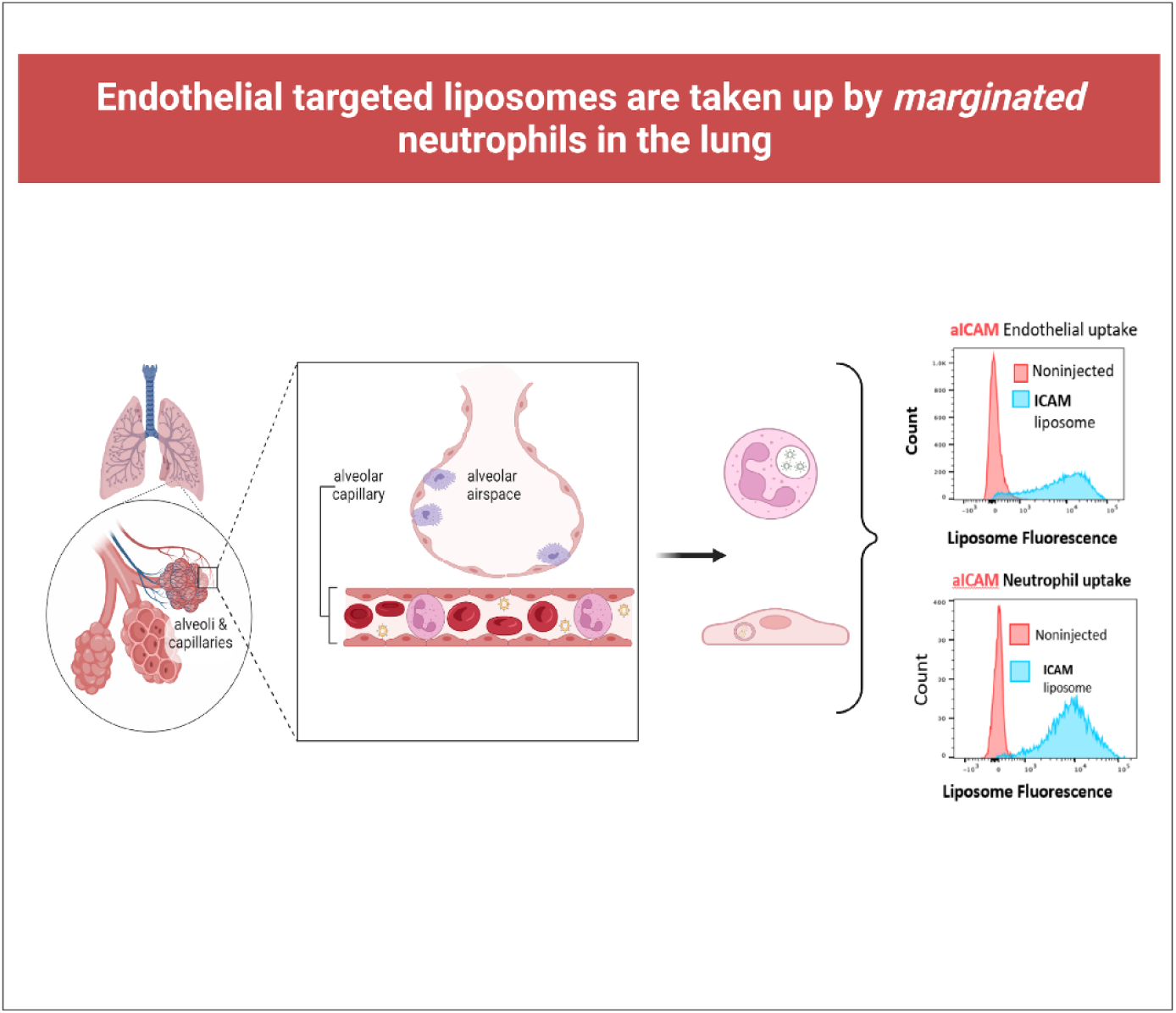

## Introduction

The decades-long goal of nanomedicine has been to localize drugs to a target organ, tissue, and/or cell type. This goal has been largely elusive. For example, in solid tumor targeting, it was shown that the top nanomedicines only achieve 0.7% of the injected dose (% ID) in the tumor, with the remainder mostly going to the main reticuloendothelial system (RES) clearance organs the liver and spleen^1,2,3^. Low organ- and cell-type specificity is a problem that continues to plague nanomedicine for various targeting methodologies aimed at diverse organs and cell types^1,2,4–7^.

A few examples of single-organ and cell-type targeting have served as a beacon for the field. More than 25 years ago, it was reported that lung endothelial cells could be targeted with nanoparticles conjugated to antibodies against endothelial surface determinants, like platelet endothelial cell adhesion molecule (PECAM), intercellular cell adhesion marker 1 (ICAM-1), and plasmalemma vesicle-associated protein (PLVAP)^4,8–10^. When injected intravenously, this endothelial-targeting method can increase small molecule drug delivery by more than 300-fold to the lungs, achieving 25% ID ^11–32^. Targeted nanoparticle delivery to the lungs thus exceeds targeted delivery to tumors and the brain, among other targets, by several orders of magnitude ^2,4^. These results have a potential clinical impact, as endothelial cells of the lungs are key players in several common diseases, such as acute respiratory distress syndrome (ARDS; the disease which kills in COVID-19), emphysema, and idiopathic pulmonary fibrosis ^33–40^. Therefore, pulmonary endothelium-targeted nanocarriers have been studied by many labs in hundreds of papers, and have achieved targeting in *ex vivo* human lungs and therapeutic effects in porcine models ^41–43^.

Several mechanisms for the high lung uptake of endothelium-targeted nanoparticles have been suggested. It was found that the first-pass advantage of the lungs (i.e., the lung capillaries are the first capillaries encountered after IV injection) only adds a small fraction (10-15% of the total) to the total nanocarrier mass delivered to the lungs ^21^. It was assumed that other phenomena led the nanocarriers to the lung: 1) Endothelial cells are more abundant in the lungs than other organs, with a surface area of 135 m^2^, maximized for gas exchange; 2) The lungs receive 100% of the cardiac output, while organs of the systemic circulation share only portions of the cardiac output; 3) Capillaries of the lungs are smaller and more tortuous than other capillary beds, potentially favoring targeted nanoparticle contact with vessel walls in the lungs over other organs ^1,22,23,44–49^.

While these studies provided much insight, these suggested mechanisms were all based on a key assumption that, until now, has yet to be rigorously tested: that in the lung, these nanocarriers were only targeting endothelial cells. This assumption was natural for multiple reasons. First, all the diverse targeting ligands (PECAM, ICAM, PLVAP, etc.) that achieve high nanoparticle lung uptake also bind to endothelial cells. Second, it was naturally assumed that endothelial cells are the only cells in capillaries that are exposed to nanocarriers in the blood (Fig 1A). What other cells in the lung could possibly have equivalent exposure to the blood? It was shown previously that nanocarriers of typical size (~100 nm) could not penetrate the < 15 nanometer inter-endothelial distance ^50^. So how could these nanocarriers possibly accumulate significantly in other lung cells?

**Figure 1:**
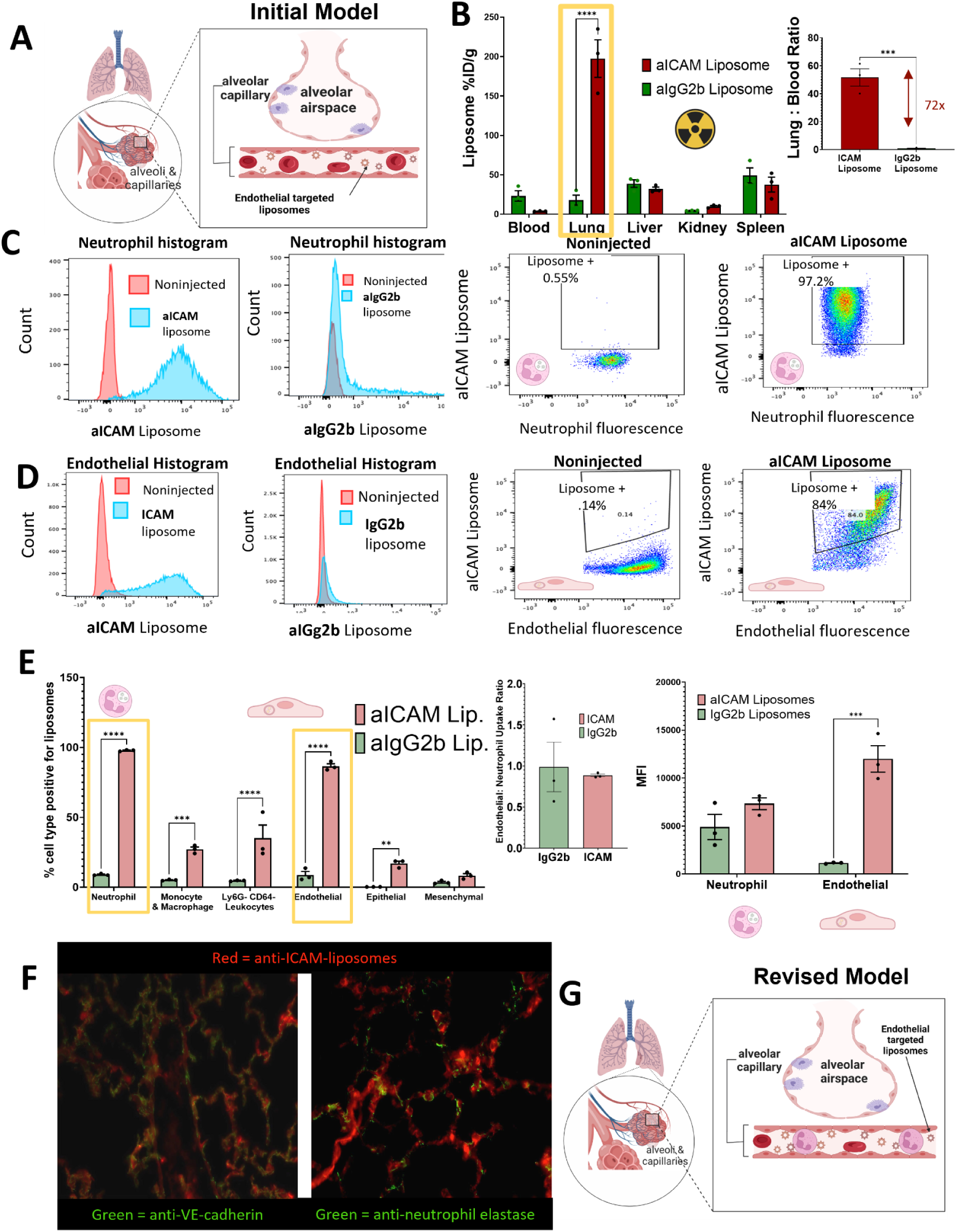
Nanoparticles that have classically been labeled as “endothelial-targeted” also target pulmonary neutrophils equally well. **A.** The classical model of lung uptake by nanoparticles conjugated to affinity ligands that bind to epitopes on endothelial cells, such as PECAM or ICAM-1. Although these epitopes are present at low concentrations on other cell types, the nanoparticles accumulate best in endothelial cells in the lungs due to the high capillary surface area, first-pass effect, and lack of competing cells in the vascular lumen. **B. Biodistribution of radiolabeled nanoparticles (liposomes) conjugated to either anti-ICAM or isotype control antibodies,** shown as percent injected dose per gram (%ID/g) 30 minutes after IV-injection into naive mice (n =3, p <0.05 by 2-way ANOVA). Inset: anti-ICAM increases the lung-to-blood concentration ratio by 72-fold compared to control (n=3, p<.05, by paired Students’ t-test. **C-E**. Flow cytometry reveals neutrophils as major cells of uptake for anti-ICAM liposomes. Mice were IV-injected with fluorescent nanoparticles similar to **B**, and 30 minutes later sacrificed to make a single-cell suspension of the lungs, which then underwent flow cytometry. **C.** Neutrophils had strong uptake of anti-ICAM (blue histogram in panel 1), but not isotype control (panel 2), liposomes; panel 4 shows similar results as a dot plot. **D.** Endothelial cells showed a similar trend as neutrophils, though with a lower % positive for liposome uptake. **E**. For each cell type, the fraction that were positive for liposome uptake, showing neutrophils as the cells with the highest percentage of uptake. Inset panel 1 shows the ratio of uptake in endothelial cells to neutrophils, while panel 2 shows the mean fluorescence intensity (MFI) of liposome-positive cells, which shows endothelial cells have only slightly higher MFIs than neutrophils. **F.** Mice treated as in C-E, but with lungs analyzed by fluorescent confocal microscopy. Red anti-ICAM liposomes are in many cell types, including endothelial cells (left panel) and within neutrophils (right panel). **G**. Revised model of uptake of nanoparticles conjugated to endothelial epitopes. An overlooked population of cells, the marginated neutrophils, resides for long periods in the pulmonary capillaries, and they take up a large fraction of the nanoparticles that accumulate in the lungs. Endothelial cells also take up such nanoparticles, but they are far from the exclusive uptake cells of the classical model. **Statistics.** All figures evaluated by 2-way ANOVA, n=3 mice, error bars are standard error of the mean (SEM). Fig 1B inset assessed by Welch’s t-test. ***= p<.001

Here, we show that an overlooked lung cell type, called marginated neutrophils, takes up these diversely targeted nanocarriers at a rate similar to or greater than endothelial cells. Marginated neutrophils reside in the capillary lumens of many organs ^51–53^, but the lungs have the most ^54,55^. A 2018 basic science study showed that, in healthy animals, there are marginated neutrophils residing in the capillaries of multiple organs^56^. They manually measured thousands of confocal images stained for neutrophils and capillary walls, which enabled definitive determination of whether neutrophils were marginated (residing in a capillary), as opposed to circulating or intra-parenchymal. They showed the <u>lungs have the highest</u> <u>concentration of marginated neutrophils, followed by the spleen, with a lower concentration in</u> <u>the liver and ski</u>n^56^.” Marginated neutrophils play a key role in patrolling for circulating pathogens, with localization to the lung being ideally suited for this function, given that the lungs have the first capillary bed encountered after a pathogen enters the bloodstream^53,57^.

Here we show that marginated neutrophils take up a large fraction of “endothelial-targeted” nanoparticles, replicating this surprising result with multiple nanomaterials, multiple targeting moieties (PECAM, ICAM, PLVAP; monoclonal antibodies, and Fab fragments), physicochemical lung targeting that does not employ targeting antibodies, in both healthy and diseased mice, and even *ex vivo* human lungs. The tendency of targeted nanoparticles to accumulate in marginated neutrophils may allow modulation of this key cell type and introduce new challenges. For example, it could decrease the payload in the target cell type, causing unwanted side effects in neutrophils; alternatively, it can open new strategies to mediate and control key side effects. We provide evidence for the mechanism by which nanoparticles are taken up in neutrophils and potential methods to limit neutrophil uptake. Since marginated neutrophils are found in other organs and diseases, we propose adding marginated neutrophils into the conceptual paradigm of the reticuloendothelial system (RES), with a special role in clearing nanoparticles that adhere to the lumenal surfaces of blood vessels.

## Methods

### Nanoparticle production and protein modification

#### Fluorescent Liposome preparation

Liposomes were prepared using thin film in round bottom glass vials. Lipids DPPC ((1,2-dipalmitoyl-sn-glycero-3-phosphocholine), cholesterol, DSPE-maleimide (1,2-distearoyl-sn-glycero-3-phosphoethanolamine-N-[maleimide(polyethylene glycol)-2000) and TopFL AF594 were combined at a molar ratio of 53.5%, 40%, 5.8% and 0.8% respectively. The thin film was then dried in a vacuum for 1hr before being resuspended in 500 microliters of phosphate-buffered saline (PBS). The mixture was then sonicated for 10 seconds three times before undergoing freeze-thaw cycles three times to improve mixing. After the freeze-thaw cycle, the solution underwent 10 extrusion cycles at 55 degrees Celsius through a membrane with 200 nm pores. After extrusion, nanoparticle size was measured using dynamic light scattering (Malvern Panalytical) to ensure liposomes were within the 150-200 nm range. PDI was less than .15 after conjugation methods detailed below and particle concentration was determined using nanoparticle tracking analysis (NanoSight, Malvern Panalytical) at a dilution of 1: 10E4 to ensure doses used for animal experiments remained at 10 mg/kg injected dose as described in Supp Fig 1.

#### Antibody conjugation

Antibodies were conjugated to liposomes by SATA-maleimide chemistry. SATA was added to the antibodies 5 times excess and incubated at room temperature for 30 minutes. To enable assessment of binding efficiency using fluorescence, the antibodies were fluorescently labeled using AF 697 fluorophores by incubating 10% of the total antibody volume for 30 minutes at room temperature in the dark. The antibodies were then purified using a desalting (Zeba) column in a centrifuge at 1530 × g for 2 minutes. Once this was completed, the antibodies were deprotected by adding 10% hydroxylamine at 34.8mg/mL and incubating at room temperature for two hours. After deprotection, the antibodies were desalted once again using a desalting column containing PBS+ 10mM EDTA and subsequently added to liposomes and incubated at 38 degrees Celsius overnight to achieve 50 antibodies per liposome as shown in Supp Fig 1. When incubation was completed, the liposome-antibody mixture was eluted through a Sepharose column. The binding efficiency of antibodies to liposomes was determined using a fluorescence plate reader. Once successful conjugation was determined, the mixture was concentrated using Amicon filters by centrifuging at 3200 × g for 2-4 hours until a volume of about 250uL was achieved. After concentrating the antibody-conjugated liposomes, the particle concentration (number of particles per mL) was measured using nanoparticle tracking analysis (Nanosight) as mentioned above.

#### Fluorescent ICAM-targeted solid lipid nanoparticle preparation

Fluorescent RNA containing lipid nanoparticles (LNPs) were prepared by preparation of an organic phase containing the following lipids: cKK-E12(Echelon Biosciences), cholesterol, 18:1 (Δ9-Cis) PE (DOPE), 1,2-distearoyl-sn-glycero-3-phosphoethanolamine-N-[azido(polyethylene glycol)-2000] (DSPE-PEG2k-azide), and AF594 PE at a lipid molar ratio of 50:37.5:9:2.5:1, respectively. All lipids are sourced from Avanti Polar Lipids except for cKKE-12 sourced from Echelon Biosciences. Scrambled siRNA (Integrated DNA Technologies) was prepared in a separate aqueous preparation at a concentration of .07 mg/mL in 50 mM citrate buffer. These mixtures are then combined in a Nanoassembler (Precision Nanosystems) using the following parameters: Flow rate ratio (FRR) of 3:1, and total flow rate (TFR) of 12 mL/min. After synthesis, LNPs are diluted 40x with 1X PBS and transferred to a 10 kDa molecular weight cutoff Amicon centrifugal filter unit (Millipore Sigma) at 3000x g for 45 min to remove excess ethanol. LNP conjugation to antibodies was carried out using DBCO-azide copper-free “click chemistry.” Azide functionalized LNPs were incubated overnight with 5x molar excess DBCO-modified antibodies at 37°C. After conjugation, LNPs were purified and sizes were measured via DLS as described above.

#### Fluorescent ICAM-conjugated polystyrene nanoparticle preparation

150 nm fluorescent carboxylate nanoparticles (Phosphorex) were exchanged into 50 mM MES buffer at pH 5.2 via a gel filtration column. N-Hydroxysulfosuccinimide (sulfo-NHS) was added to the particles at 0.275 mg/mL, before incubation for 3 minutes at room temperature. EDCI was added to the particles at 0.1 mg/mL, before incubation for 15 minutes at room temperature. ICAM was added to the particle mixture at 50 ICAM per nanoparticle, before incubation for 3 hours at room temperature while vortexing. Beads were then purified, and sizes were measured via DLS, as described above.

#### Radiolabeling conjugated liposomes

Using the above mentioned technique, nonfluorescent liposomes were synthesized and conjugated with ICAM, PECAM, PLVAP, or IgG2b antibodies. Non-specific IgG2b antibodies were labeled with I-125 and conjugated onto liposomes alongside the above antibodies at 2% radiolabeled IgG2b + 98% unlabeled antibody to label the liposomes. I-125 labeling was achieved by using Pierce Iodogen radiolabeling reagent and purifying with desalting (Zeba) columns. To confirm the radiochemical purity of the antibody, thin-layer chromatography was used.

#### Variation of nanoparticle size and charge

To maintain a consistent formulation and vary the size and charge of our liposomes, they were formulated using the microfluidic mixing method, using a NanoAssmblr Ignite.

An organic phase containing a mixture of lipids (54% DPPC, 39.9% Cholesterol, 5.8% DSPE-PEG2k-Maleimide, 0.3% 18:1 PE-TopFluor® AF594 by moles) dissolved in ethanol with total lipid mix concentration of 20 mM was mixed with an aqueous phase (1x PBS) at various flow rate ratios and total flow rates. Liposomes were dialyzed against 1x PBS in a 10kDa molecular weight cut-off cassette for 2 h, sterilized through a 0.22 μm filter, and stored at 4 °C.

To produce 200 nm liposomes, we set the NanoAssemblr Ignite’s flow rate to 0.5 mL/min with an aqueous-to-lipid ratio of 0.5:1 (a heating attachment was used to keep the lipid mix temperature at 52 °C). To produce 150 nm liposomes, we set the flow rate to 1 mL/min with an aqueous-to-lipid ratio of 3:1. To produce 70 nm liposomes, we set the flow rate to 6 mL/min with an aqueous-to-lipid ratio of 3:1.

100 nm DOTAP liposomes were formulated using a total flow rate of 0.5 mL/min and an aqueous-to-lipid ratio of 1:1 (50% DOTAP, 24.86% DPPC, 19.04% Cholesterol, 5.8% DMG-PEG2K, 0.3% 18:1 PE-TopFluor® AF594).

70, 150, and 200 nm liposomes were conjugated with aPECAM antibodies at a coating density of 50 mAb/liposomes using SATA-maleimide chemistry. Conjugation efficiency was validated using size exclusion chromatography as previously described. Size and zeta potential are described in Supp Fig 9 for 70, 150, 200, and 110 nm DOTAP liposomes.

### Microscopy

#### Thick-cut 3D histology

30 minutes after injection with AF594 labeled liposomes, C57/BL6 mice were sacrificed and their lungs were perfused with cold PBS. After perfusion, the lungs were then inflated using 0.8mL of 2% paraformaldehyde and allowed to rest in the same fixative overnight. The next day, the fixed lungs were rinsed three times with ice-cold 1x PBS, then stored at 4 C until ready to be cut. Samples were then positioned, cut into thick sections, and covered in low melting agarose, allowing them to cool before sectioning. Samples are then sectioned on a vibratome. After sectioning, samples are prepared for staining using a permeabilization/wash buffer (1xPBS + 1% Triton X-100 for 1 hour at room temperature. Samples are then incubated in 1ml Staining Buffer (PBS+1% BSA+0.3% Tritonx100) and desired primary antibodies for 48 hours at 4C. Samples are then washed three times with a wash buffer at intervals of 30 minutes at room temperature. After this, mice were stained with 1 mL staining buffer and incubated for 24-48 hours at 4 C. Samples were washed in Perm/Wash Buffer three times for 30 minutes at room temperature. After this wash, samples were incubated in 2mL Scale A2 (4M Urea+10% Glycerol+0.1% Triton X-100 in water) for 4-7 days. This was followed by 2 mL of Scale B4 (8M Urea+0.1% Triton X-100 in water) and a 1:2000 dilution of DAPI overnight at room temperature. After DAPI incubation, fresh Scale B4 was added and allowed to incubate for 24-48 hours at room temperature, then transferred to Scale A2 at 4C for 2 to 10 days before imaging.

#### Confocal microscopy on lung and spleen after cecal ligation and puncture (CLP model)

2 hours after cecal ligation and puncture (CLP) injury described below, mice were injected with fluorescently labeled LNPs. 30 minutes later, Alexa fluor-647 labeled Ly6G was injected and circulated for 5 minutes before the animals were sacrificed. The lungs and spleen were harvested and frozen after gravity perfusion at 25 cm with cold PBS. 10 um tissue slices were cryosectioned for imaging using Leica TCS SP8 confocal microscopy.

#### Intravital microscopy for visualizing pulmonary capillaries

Mice were sedated with intraperitoneal pentobarbital. The right jugular vein was cannulated, and fluorescent LY6G and CD31 were injected. Mice were then intubated and placed on a ventilator (Harvard Apparatus). Mice were ventilated with a tidal volume of 10uL of compressed air per gram of mice weight, a respiratory rate of 130-140 breaths per minute, and positive-end expiratory pressure of 2 cm H2O. A left thoracotomy was performed on the mouse in a lateral decubitus position and a thoracic window was applied and secured to the stage. 25 mm Hg of suction was applied using a vacuum regulator (Amvex Corporation) to immobilze the lung. Mice were placed on a microscope, and endothelium and marginated neutrophils were visualized. Fluorescent liposomes were injected, and video recordings via microscope were collected at 2 minutes, 15 minutes, and 30 minutes after injection. After collecting recordings, mice were euthanized.

### Animal Studies

#### Animal Study protocols

All animal studies were performed strictly per the Guide for the Care and Use of Laboratory Animals as adopted by the National Institute of Health and approved by the University of Pennsylvania and the Children’s Hospital of Philadelphia’s Institutional Animal Care and Use Committee. All animal experiments used male C57BL/6 mice at 6-8 weeks old (Jackson Laboratories). The mice were maintained at 22-26°C and adhered to a 12/12h dark/light cycle with food and water ad libitum

#### Nebulized lipopolysaccharide (LPS) Model

Mice were exposed to nebulized LPS in a ‘whole-body’ exposure chamber, with separate compartments for each mouse (MPC-3 AERO; Braintree Scientific, Inc.; Braintree MA). To maintain adequate hydration, mice were injected with 1mL of sterile saline, 37C, intraperitoneally, immediately before exposure to LPS. LPS (L2630-100mg, Sigma Aldrich) was reconstituted in PBS to 10mg/mL and stored at −80C until use. Immediately before nebulization, LPS was thawed and diluted to 5 mg/mL with PBS. LPS was aerosolized via a mesh nebulizer (Aerogen, Kent Scientific) connected to the exposure chamber (NEB-MED H, Braintree Scientific, Inc.). 5 mL of 5mg/mL LPS was used to induce the injury. Nebulization was performed until all liquid was nebulized (~20 minutes).

#### Complete Freund’s Adjuvant Footpad inflammation model

Complete Freund’s adjuvant (CFA) model is commonly used to elicit local inflammation ^58^. Briefly, mice were anesthetized with 2% (vol/vol) isoflurane, and the left hind paw was sterilized with a 70% alcohol pad. Then, 20μL of CFA emulsion (Sigma-Aldrich) was subcutaneously injected into the central plantar region of the left hind paw. After 24 hours, animal paws and organs were harvested for various assays and evaluations.

#### Cecal Ligation and Puncture (CLP) model of sepsis

Cecal ligation and puncture (CLP) is considered the gold-standard model in sepsis research, in which a systemic inflammatory response is triggered by the polymicrobial infection in the abdominal cavity and the following blood bacterial translocation ^59^. Briefly, mice were anesthetized with intraperitoneal injection of the mixture of ketamine (80 mg/Kg body weight) and xylazine (7 mg/Kg body weight). Once mice were fully anesthetized, they were placed supine for surgery. Firstly, a 1 cm midline skin incision was made with a scalpel and subsequently, the cecum was identified and exposed from the peritoneal cavity using blunt anatomical forceps.]. Once the cecum was exposed, 25-30% of the cecum was ligated to achieve a mild-grade sepsis. Following ligation, a 21 gauge needle was used to perforate the cecum by a through-to-through puncture near the ligation site while avoiding the mesenteric blood vessels.

After the puncture, the needle was removed, and the ligated cecum was gently squeezed with a small amount of feces extrusion and then the cecum was placed back into the peritoneal cavity. The abdomen was then closed using a silk surgical suture 4-0 with a cutting needle. Then, the punctured cecum was gently located back into the peritoneal cavity, and the abdomen was closed by using the silk surgical suture 4-0 with a cutting needle. Postoperatively, the mice were resuscitated with subcutaneously injected normal saline at 1ml per 25g and placed under a heat lamp until they were fully recovered. Mice were then placed back into cages with convenient access to water and food. Based on the experiment design, 2 hours after surgery, animals were injected with CAM-targeted nanoparticles and then used for various assays and evaluations.

#### Biodistribution Measurement

For biodistribution, naïve mice or nebulized lipopolysaccharide (neb-LPS) treated mice were injected intravenously with liposomes at a 10mg/kg lipid dose. For neb-LPS treated groups, mice were nebulized four hours before injection of nanocarrier as described above. At 10 minutes, 30 minutes, or 1 hour after injection with liposomes, mice were euthanized. Blood, lungs, liver, heart, kidney, spleen, injection site, and syringe were collected for analysis. Based on a known injected dose of radioactivity, quantified by Wallac 2470 Wizard gamma counter (PerkinElmer Life and Analytical Sciences-Wallac Oy, Turku, Finland), this was used to determine the biodistribution by calculating the percent injected dose per gram of tissue.

An in vivo imaging system (IVIS) was used to analyze biodistribution. Liposomes were prepared by thin film hydration using the lipid formulation described earlier. Liposomes were formed by hydrating the lipid film with 500uL of a 10ug/mL solution of indocyanine green dissolved in 1X PBS and extruded as described above. Liposomes were then covalently conjugated to either IgG or anti-PECAM antibody (MEC13) to achieve 50 mAb/liposomes utilizing SATA-maleimide chemistry, as described above. Following size exclusion chromatography by Sepharose column to remove unbound drug and antibody, liposomes were injected into mice at 10 mg/kg and circulated for 30 minutes. Mice were then sedated under isoflurane and imaged *in vivo* utilizing IVIS. After euthanization, organs were harvested and imaged utilizing an Ex/Em of 720/800 on IVIS.

#### Cell type distribution and flow cytometry

C57/BL6 mice were injected with liposomes and euthanized after 30 minutes (n=3). Lungs were collected and triturated before incubating in a digestive solution of 2 mg/mL collagenase and 100uL of 2.5mg/mL DNase for 45 minutes, to prepare a single-cell suspension. After incubation, the cells were strained through a 70uL cell strainer and washed with PBS. The supernatant was discarded and ACK lysis buffer was added for 5 minutes on ice to lyse any remaining red blood cells. After cell lysis, the cells were washed again at 1224 rpm for 5 minutes at 4 degrees Celsius and then counted using an automated cell counter (Countess, Thermo Fisher) to obtain the cell concentrations. The goal was to achieve a concentration of 1 × 10^6^ cells/mL. After cell counting, the suspension was washed with FACS buffer and incubated with Fc block for 15 minutes, followed by 30-minute antibody incubation using CD45, CD64, and Ly6G for the leukocyte panel and CD31 and EPCAM for the endothelial cell panel. Fluorophores and markers used for the identification of cell types as well as the gating strategy are summarized in Supp. Fig. 2. After antibody incubation, the cells were fixed with 4% PFA before analysis on a flow cytometer (LSR Fortessa, BD BioSciences).

#### Validation of nanoparticle crossover index (NCI)

To validate the presence of artifacts caused by single-cell preparation of lungs for flow cytometry, one set of C57/BL6 mice was injected with 10 mg/kg of Alexa-594-labeled liposomes, while another set of mice received the same dose of Alexa-488 labeled liposomes (and route of delivery). The left lungs of each mouse were combined into a single tube for processing for flow cytometry (the right lungs were treated similarly, as a replicate), the combined tissues were then processed for flow cytometry as described above. As controls, mice injected with only Alexa-594 or Alexa-488 labeled CAM-targeted liposomes were also prepared for flow cytometry to establish baselines.

#### Identification of neutrophil compartments

To distinguish the different neutrophil compartments found in the lung, we developed the following flow cytometry protocol to identify the following compartments: neutrophils found in the airspace, interstitial, and marginated neutrophils. We injected 10 mg/kg aPECAM liposomes labeled with AF594 and allowed them to circulate for 30 min. 5 min before sacrifice, mice were injected with RED (AF700) anti-Ly6G labeled antibody. Mice were then sacrificed, and blood was pulled from the inferior vena cava (IVC) using a 25 gauge needle pre-coated with .5 M EDTA. The IVC was then cut and the mouse was prepared for bronchoalveolar lavage (BAL). To prepare for BAL, the salivary glands were dissected to expose the strap muscles around the trachea. After exposing the trachea, an incision is made and a catheter is inserted and tied in place with a suture. Lungs were then lavaged with three washes of .8mL of 1x PBS with 0.5 mM EDTA and stained for later analysis. After BAL washes, lungs were then disaggregated to generate single-cell suspensions and stained for flow cytometry using the flow cytometry protocol described above. Resultant single cell preparation from lung tissue is later post-stained with VIOLET (AF 711) anti-Ly6G labeled antibody. This results in the following: blood samples from the IVC represent circulating neutrophils stained only RED, marginated neutrophils become stained with **both** RED and VIOLET, interstitial neutrophils stained only with VIOLET, and BAL samples are stained directly with VIOLET. We then stained for CXCR4 and CD11b presentation for additional surface marker studies.

#### Complement Studies In vitro

IgG, aPECAM, and aICAM liposomes were generated as previously described. On the day of the experiment, fresh serum from mice was obtained. 20uL of fresh serum was incubated 20 uL of either IgG, aPECAM, or aICAM liposome each at a concentration of 2 × 10^12^ liposomes per mL (determined beforehand via nanoparticle tracking analysis [NTA] in filtered 1xPBS). Alexa Fluor 488 labeled C3 was added at a concentration of 1.2 mg/mL to match the endogenous quantity of C3 in serum. Everything was then incubated at room temperature for 20 minutes. After this time, the resulting 40uL of liposome-serum mixture is diluted in 960 uL of filtered PBS to quench C3 opsonization. A further 20uL was diluted in 980uL of filtered PBS to assess via NTA.

#### In vivo

As previously described, fluorescent aPECAM liposomes were generated. They were administered to wild-type (C57/B6) or C3 knock-out mice (derived from C57/B6) at 10 mg/kg. After a 30-minute circulation period, the lungs were perfused and processed into a single-cell suspension. The fluorophores and markers used to identify cell types and the gating strategy are summarized in **Supp. Fig. 2**. After antibody incubation, the cells were fixed with 4% PFA before analysis on a flow cytometer (LSR Fortessa, BD BioSciences).

#### C3a ELISA

ELISA was performed to assess activated C3a levels *in vitro*, per manufacturer protocol. For *in vitro* measurement, 20 µL of fresh serum was incubated with 20 µL of either bare (no antibody, but still PEGylated), IgG, aPECAM, or aICAM liposomes (2 × 10^12^ liposomes/mL) for 15 min, EDTA was added to a final concentration of 20 mM, to inhibit further complement activation.

### Nanoparticle administration in human lungs

Human lungs used in this study were from de-identified non-used lungs donated for organ transplantation following an established protocol (PROPEL, approved by the University of Pennsylvania Institutional Review Board) with informed consent in accordance with institutional and NIH procedures. Consent was provided by next of kin or healthcare proxy. The institutional review board of the University of Pennsylvania approved this study, and all patient information was removed before use. While this tissue collection protocol does not meet the current NIH definition of human subject research, all institutional procedures required for human subject research at the University of Pennsylvania were followed throughout the reported experiments. Lungs were kept at 4°C and were used within 12 hours of procurement. Lungs were inflated to prevent atelectasis and allow for adequate perfusion. The pulmonary artery was cannulated and then perfusate-containing tissue dye was infused to allow identification of the perfused portions of the lungs. Using nanoparticle synthesis methods as described above, human anti-ICAM antibodies were conjugated to liposomes. The dose for human injection was scaled up to 40 mg/kg lipid dose, based on the mass of the donated lung. Following perfusion with acellular albumin-based perfusate for 10 minutes, 3ml of anti-ICAM liposomes were infused and left for 10 minutes followed by perfusion. Subsequently, 4g of perfused tissue was collected and processed to single-cell suspension for flow cytometry similar to the mouse tissue as above. Flow cytometry markers and fluorophores used as well as the gating strategy used to identify different cell populations can be found in Supp Figs 2,3, and 5 and summarized in Supp Tables 3-6.

### Statistics and Software

All statistics and software were analyzed using GraphPad Prism. Tests used are indicated in the figure legends.

## Results

### Nanoparticles classically thought to target the lungs’ endothelial cells target marginated neutrophils just as strongly *in vivo*

Our initial goal was determining what lung cell types take up endothelial-targeted nanoparticles. As the field implicitly thought the answer would be almost exclusively endothelial cells as shown in **Figure 1A**, we set out to more rigorously test this hypothesis. We noted that some “endothelial epitopes,” such as ICAM, are known to be expressed on leukocytes and endothelial cells, though at a lower concentration ^50, 72,73^. Therefore, we constructed nanoparticles conjugated to anti-ICAM antibodies to test cell type specificity more rigorously. As our nanomaterial, we employed liposomes, since they are by far the most clinically employed nanoparticles ^60^. As shown in **Figure 1B**, radiolabeled anti-ICAM liposomes strongly accumulated in the lungs of naive mice 30 minutes after IV injection, with a lung-to-blood ratio 72x higher than that for liposomes conjugated to untargeted IgG2b isotype-control antibodies. This biodistribution closely resembles those reported many times by our group and others ^11–29^, confirming that the function of “endothelial-targeted” nanoparticles is similar to that of prior studies.

Flow cytometry is the most quantitative method to investigate which lung cell types take up the nanoparticles. In this method, fluorescently labeled nanoparticles are injected IV. Then the lungs are disaggregated into single-cell suspensions, stained with cell-type-specific markers, and quantified in a flow cytometer. Here, anti-ICAM liposomes containing an Alexa Fluor 594-conjugated lipid were injected IV and 30 minutes after, lungs were disaggregated and analyzed using the flow cytometry gating strategy described in **Supp Fig 3**. Neutrophils had very strong uptake of anti-ICAM liposomes compared to untargeted control (**Figure 1C**) with 97.2% of neutrophils positive for anti-ICAM liposomes. Endothelial cells also showed strong uptake of ICAM-targeted liposomes; however, the percentage of positive cells was surprisingly less than neutrophils, at 84% (**Figure 1D**). When compared amongst other cell types, neutrophils and endothelial cells had the most uptake of ICAM-targeted liposomes compared to untargeted IgG2b-conjugated liposomes (**Figure 1E, panel 1**). Interestingly the endothelial-to-neutrophil uptake ratio was similar between anti-ICAM and isotype-control liposomes (**Figure 1E Panel 2**). However, the mean fluorescence intensity (MFI) of anti-ICAM liposomes was much higher in endothelial cells than in neutrophils (**Figure 1E Panel 3**). Also, we found multiple cell types that were positive for liposome fluorescence, but the total uptake was small compared to neutrophils (**Fig 1E, panel 1**). Thus, the core new finding is that “endothelial-targeted” nanoparticles (here anti-ICAM liposomes) do not solely target the endothelial cells; instead, neutrophils and endothelial cells seem to be nearly co-equal in their nanoparticle uptake.

This finding is also supported by additional studies looking at the total distribution of aICAM-conjugated liposome-positive cells; i.e., gating first to identify all liposome-positive cells, followed by analyzing the fractions of liposome-positive cells represented by each cell type, as shown in **Supp Fig 4**. Gating this way reveals that of the total liposome-positive population, 40% were endothelial cells 40% were neutrophils, while an additional 20% were leukocytes of neither macrophage nor neutrophil lineage. This means that 60% of “endothelial-targeted liposomes” are not being taken up by endothelial cells and are instead being taken up by innate immune cells, with neutrophils taking the lion’s share among immune cells, and at a co-equal amount to endothelial cells **Supp Fig 4**.

To confirm this surprising flow cytometry result, we used confocal microscopy to visualize the distribution of nanoparticles in lung tissue directly. Anti-ICAM liposomes can be seen in endothelial cells and neutrophils (**Figure 1F**). This discovery of strong neutrophil uptake in “endothelial-targeted” nanoparticles was surprising because the endothelial cell is usually depicted as the only cells facing the blood in capillaries. However, marginated neutrophils reside in the capillaries (especially of the lungs) for long periods, where they are perfectly situated to take up nanoparticles ^51–53^. Thus, we revised our schema to reflect the paradigm shift in our proposed mechanism of “endothelial-targeted” nanoparticle uptake (**Figure 1G**).

### Neutrophil uptake is generalizable to multiple targeting moieties that are classically described as targeting endothelial cells only

Having shown that anti-ICAM nanoparticles are taken up strongly by marginated neutrophils in the lungs, we wanted to know if this result generalized to other “endothelial-targeted” nanoparticles. Several proteins expressed on endothelial cells have been used as targets to enrich lung uptake of nanoparticles (**Figure 2A**): ICAM, PECAM (which is not closely related genetically to ICAM), PLVAP, thrombomodulin, APP-1, and more. Therefore, we conjugated liposomes to monoclonal antibodies targeting (separately) ICAM, PECAM, and PLVAP, three of the most studied endothelial surface markers. As shown in **Figure 2B**, all three of these liposomes accumulated in the lungs, to extents similar to those previously reported ^9–17,20,29,61–64^. As with previous studies, these high lung uptakes are achieved by 10 minutes of circulation time and remain constant for at least half an hour (**Figure 2C**).

**Figure 2:**
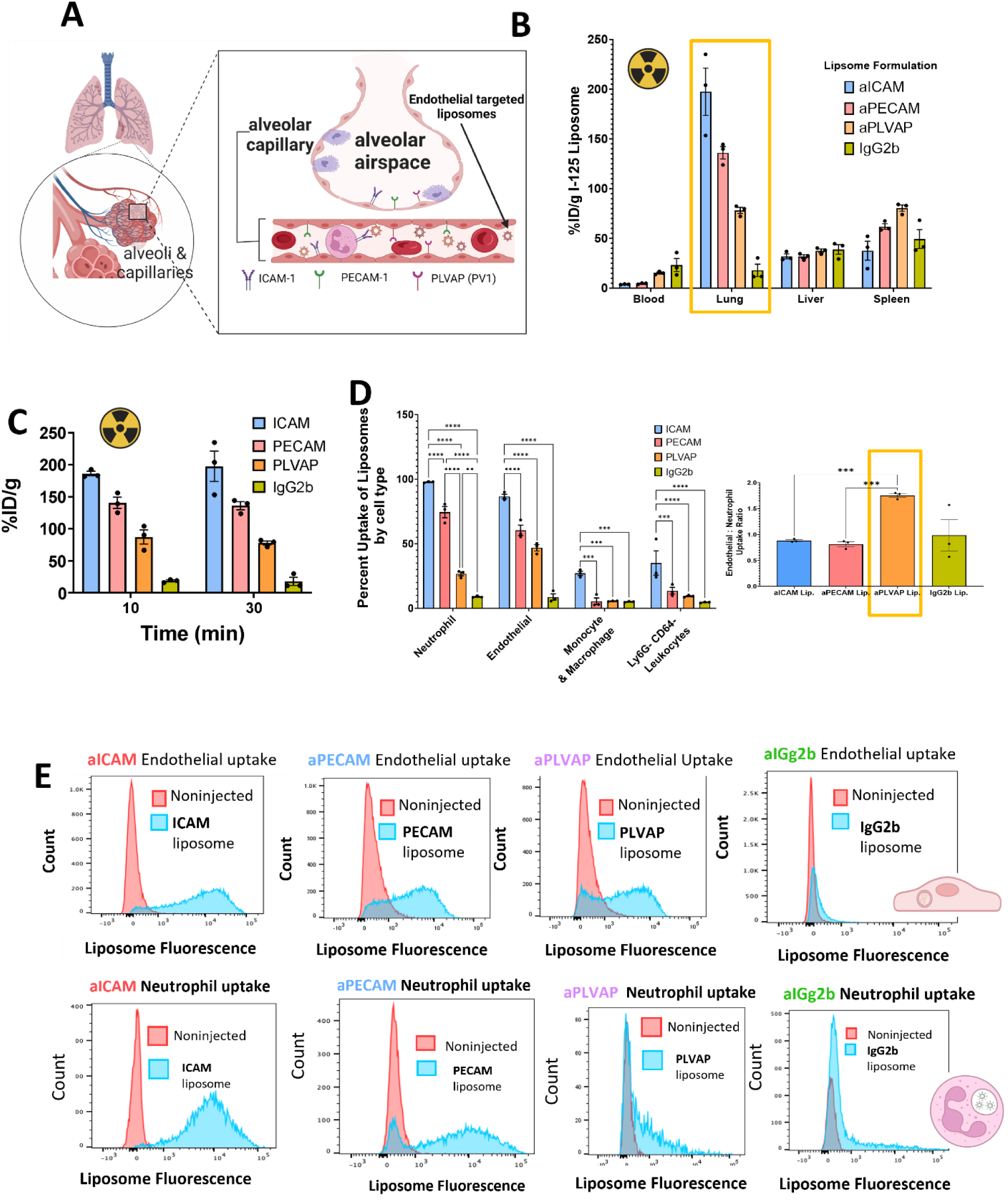
Neutrophil uptake is generalizable to multiple targeting moieties classically described as targeting endothelial cells only. **A.** Schema representing the known distribution of common epitopes used to target the pulmonary endothelium. **B.** Biodistribution 30 minutes after IV injection of radiolabeled liposomes conjugated to antibodies that bind to putative endothelial targets: ICAM, PECAM, PLVAP, and isotype control. Displayed as percent injected dose per gram (%ID/g), the lungs are the dominant target organ). **C.** Nanoparticle biodistribution to the lungs is fast, peaking before 10 minutes and remaining stable up to 30 minutes. **D.** Cellular distribution of liposome uptake (measured by flow cytometry) among all targeting moieties. The two most commonly used moieties to target the pulmonary endothelium, anti-ICAM and -PECAM, both have strong neutrophil uptake. PLVAP, less commonly used for this purpose, has lower overall lung uptake due to lower access, but a higher endothelial-to-neutrophil targeting ratio (inset). **E.** Representative histograms from flow cytometry performed in D, showing specific neutrophil uptake in all three targeting moieties but not isotype control. **Statistics** Fig 2D, 2-way ANOVA with Tukey post-hoc test **=p<0.01, ***=p<.0002, ****=p<.0001. Fig 2D inset, Brown-Forsythe and Welch ANOVA Test using Dunnet’s T3 post hoc test***=p<0.01

Using these liposomes uniquely targeted to specific endothelial epitopes, we injected them into mice and performed flow cytometry, as with anti-ICAM liposomes in **Figure 1**. We found that there is significant neutrophil uptake for all three types of targeted liposomes (**Figures 2D and E**). The endothelial-to-neutrophil uptake ratio was nearly identical for ICAM and PECAM (both less than 1, meaning neutrophils favored). Still, PLVAP had an endothelial-to-neutrophil uptake ratio greater than 2x higher. Thus, strong uptake by marginated neutrophils is likely common to all “endothelial-targeted” nanoparticles. This is interesting considering that while neutrophils are known to express ICAM and PECAM on their surface^24,65,66^, endothelial cells have far more of each present on their surface. However, some targets, such as PLVAP, which is uniquely expressed on endothelial cells, yield more endothelial-specific nanoparticle uptake and thus may be preferred in applications when neutrophil avoidance is particularly important.

As with ICAM-liposomes above, we once again analyzed the data using the opposite gating strategy: gate first on liposome-positive cells, then examine which fraction of liposome-positive cells are comprised by each cell type. For both anti-ICAM and -PECAM liposomes, among liposome-positive cells, ~40% were neutrophils, ~40% were endothelial cells, and the remaining 20% were from other immune cells (**Supp Fig 4**). Additionally, endothelial cells heavily prefer aPLVAP-conjugated liposomes with 80% of liposome-positive cells being endothelial cells.

### The majority of CAM-targeted liposomes are taken up by lung-marginated neutrophils

Having shown that uptake of endothelial-targeted liposomes by neutrophils is generalizable to multiple targeting moieties, we next wanted to determine in which neutrophil compartment in the lung our CAM-targeted nanocarriers were being taken up. We hypothesized that the lung-marginated pool would most likely have our nanocarriers in this compartment as they are primed for their key role in the surveillance and patrol of the lung for circulating pathogens ^56,67^.

Evaluating data for potential artifacts is important for any result contradicting a commonly held assumption. Therefore we set out to confirm that our nanoparticles were taken up by neutrophils *in vivo*, rather than the neutrophils taking up nanoparticles artifactually during *ex vivo* preparation of the single-cell suspension made for flow cytometry. During the preparation of single-cell suspensions for flow cytometry, there is potential for loosely associated nanoparticles to dissociate and be taken up by other cells *ex vivo*, which we define as “nanoparticle crossover.” To ensure that this nanoparticle crossover between cell types did not significantly affect the results, we developed a novel, generalizable protocol to quantify the degree of this crossover as shown in the schematic in **Figure 3A** and detailed in our methods section. In brief, one mouse was injected with Alexa 594 liposomes, while a second mouse was injected with Alexa 488 liposomes. The left lungs of each mouse were put together into a single tube, and then the single-cell-dissociation and flow cytometry were performed on that tube. If nanoparticle crossover happens at a high rate, we should be able to observe cells that are positive for <u>both</u> Alexa-594 and -488. As shown in **Figure 3B**, we found that only a small fraction, less than 7% in lung endothelial cells and less than 10% in neutrophils were double positive for Alexa-594 and -488, suggesting that a small fraction of liposomes are indeed able to crossover from one lung to the other during the *ex vivo* preparation. Even among double-positive cells, the mean fluorescence of the “second fluorophore” was consistently low, indicating low levels of nanoparticle crossover. Further, some cross-over fluorescence is likely to be fluorescence bleed-through between the two fluorescent filters. Therefore, the quantity of nanoparticle crossover artifacts is insufficient to greatly impact the bottom-line results that neutrophils and endothelial cells both strongly uptake ICAM-targeted nanoparticles in the lungs. We further define crossover capability as nanoparticle crossover index (NCI). It is defined as the double positive cell fraction divided by the sum of the double positive fractions, Alexa-488 only, and Alexa-594 positive. Thus an NCI value of 1 would indicate complete *ex vivo* mixing of the nanoparticles in a preparation. **Figure 3C** shows the NCI is very low (<< 1) for both endothelial cells and neutrophils. To ensure we can detect artifactual uptake of nanoparticles during the single-cell preparation process, we used as our positive control intratracheal instillation of liposomes (as opposed to our IV injection of anti-PECAM nanoparticles). We recently showed that after intratracheal instillation of liposomes into the lungs, the liposomes remain in the airspace of the lungs, not taken up cells, for many hours^68^; ~80% of liposome remain extracellular, in the alveolar airspace, at 4 hours after liposome instillation. As hypothesized, after intratracheal instillation, liposomes had a very high nanoparticle cross-over index (NCI) of 51.2% (**Supp Fig 5**), showing that the apparent marginated neutrophil uptake mostly occurred *ex vivo*, rather than in vivo. By contrast, IV injection of anti-PECAM nanoparticles had a very low NCI, showing that the vast majority of marginated neutrophil uptake of these nanoparticles did occur *in vivo* (**Supp Fig 5**). Thus, it is clear that our flow cytometry results showing strong neutrophil uptake of nanoparticles represents an in vivo event, and is not artefactual.

**Figure 3.**
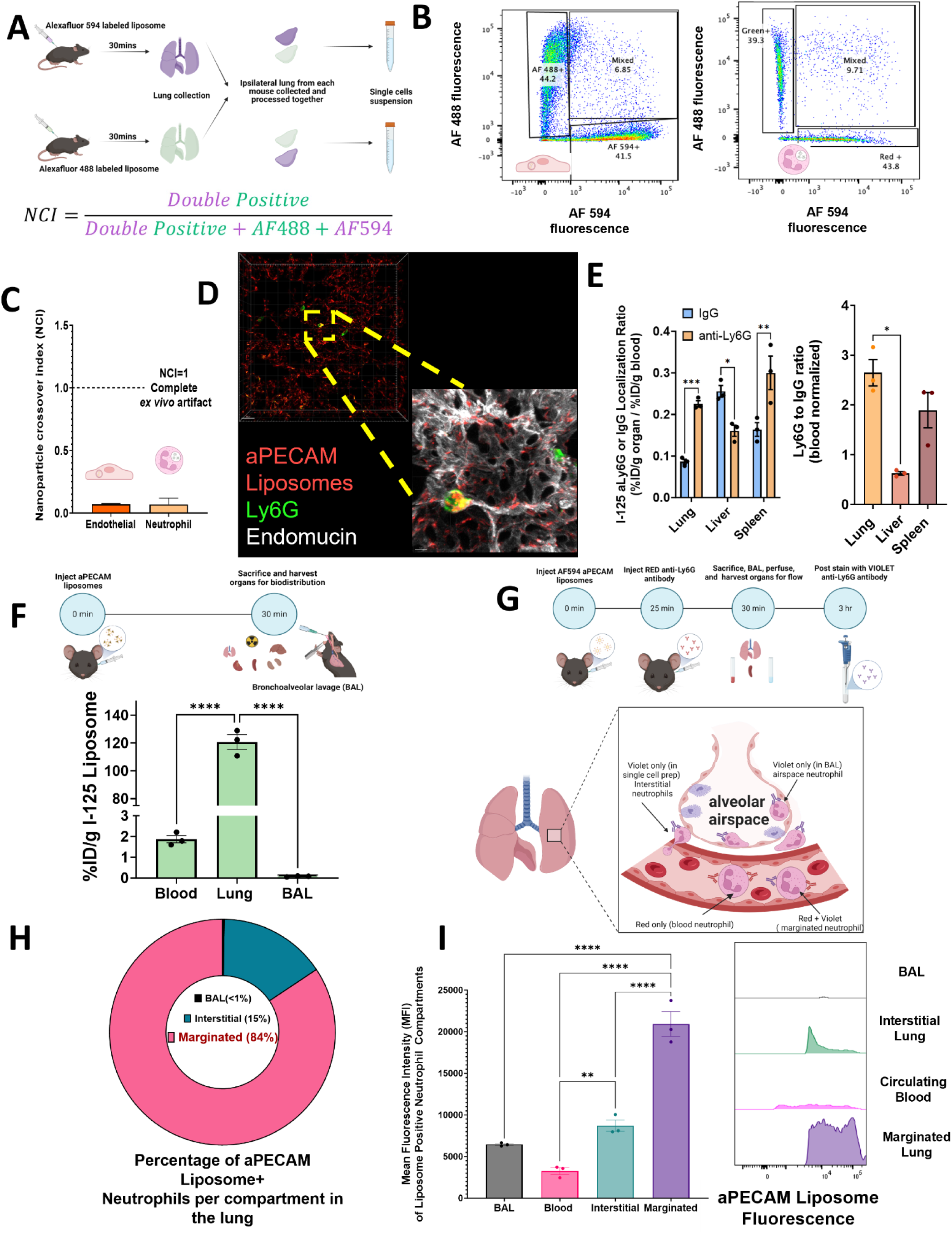
Most CAM-targeted liposomes are taken up by marginated neutrophils in the lungs. If a nanoparticle localizes to the lungs but is not internalized into cells, it is possible that it could be taken up into cells after the mouse is sacrificed, especially during the single-cell-suspension step before flow cytometry. This could artificially inflate the fraction of a cell type taking up the particle. To quantify this potential artifact, we devised the following generalizable protocol (**A**): One mouse was given Alexa-594-labeled liposomes, and another had the same liposomes (and delivery route), labeled with Alexa-488. The left lungs of each mouse were combined into a single tube for processing for flow cytometry (the right lungs were treated similarly, as a replicate). **B**. Flow cytometry from an example mouse. The vast majority of endothelial cells were positive for only one color liposome (594 or 488, in the bottom-right or top-left quadrants, respectively). There were 6.85% of endothelial cells that were positive for both liposome colors, suggesting that a small fraction of liposomes can pass from one lung to the cells of the other while in the post-mortem single-cell suspension. This is similar for neutrophils from the same experiment with about 9.81% of neutrophils being mixed **C**. We define the nanoparticle crossover index as listed in the figure, and it gives the fraction of liposome-positive cells that received a significant fraction of their liposomes post-mortem in the single-cell suspension. We show that this value for endothelial cells and neutrophils is very low. For comparison an NCI =1 would indicate complete *ex vivo* transfer of our nanoparticle following flow cytometry preparation. **D.** We used high-resolution microscopy to visualize neutrophils in the capillary lumen taking up our CAM-targeted liposomes **E.** To quantify marginated neutrophils in each organ, we IV-injected radiolabeled anti-Ly6G, 5 minutes later sacrificed, and perfused. After normalizing each organ’s radioactivity to blood radioactivity (left panel), it is evident that marginated neutrophils are higher in the lung than the liver, especially when compared to control radiolabeled (non-avid) IgG. In the right panel, we normalized the anti-Ly6G signal by control IgG, which shows the lungs have ~4x higher concentration of marginated neutrophils than the liver. **F.** To determine if endothelial-targeted nanocarriers were leaking out of the endothelium and into the alveolar airspace, we IV-injected aPECAM-liposomes, sacrificed 30 minutes later, perfused the lungs, and then measured radioactivity in the airspace (via bronchoalveolar lavage [BAL]), and the residual lungs and blood. The lungs have by far the highest uptake, showing nearly no leak of nanoparticles into the airspace. **G.** We next developed a flow cytometry protocol to distinguish the different neutrophil compartments in the lung: neutrophils in the airspace, interstitial, and marginated neutrophils. Briefly, we inject liposomes labeled with AF594 and allow them to circulate for 30 min. 5 min before sacrifice, mice were injected with RED anti-Ly6G antibody. Mice were then sacrificed and blood and bronchoalveolar lavage (BAL) fluid samples were drawn for later analysis. Resultant single cell preparation from lung tissue is later post-stained with VIOLET anti-Ly6G antibody. This results in the following: blood samples representing circulating neutrophils stained only RED, marginated neutrophils stained with **both** RED and VIOLET, interstitial neutrophils stained only with VIOLET, and BAL samples stained only VIOLET. **H.** Gating on liposome-positive cells first, then identifying the percentage of each compartment, we find that in the lung, marginated neutrophils account for 84% of total liposome-positive neutrophils. **I.** MFI and histogram show that among neutrophils, marginated neutrophils in the lungs take up more nanoparticles than those in the blood, interstitium, or airspace. **Statistics**. n=3 per group unless otherwise stated. 1-way ANOVA with Tukey post hoc test *p<0.05, **p<0.01,***p<0.001, ****p<0.0001

Having validated that nanoparticle uptake by lung neutrophils is accurate, we next endeavored to show that such neutrophils in the lungs are indeed marginated (residing in the intravascular space). First, we performed 3D histology on thick, cut lung tissue sections. As shown in **Figure 3D** and **Supplemental Movie 1**, we observed that the neutrophils taking up our nanocarriers are indeed intravascular (inside the lumen of capillaries).

Next, we quantified marginated neutrophils in the lungs compared to other organs, by radiolabeling marginated neutrophils *in vivo.* We intravenously (IV) injected radiolabeled anti-Ly6G antibodies (specific to neutrophils), and then 5 minutes later measured the amount of radioactivity in each organ. The injection of anti-Ly6G only 5 minutes before sacrifice prevents antibody extravasation into parenchyma^69^. Here, we radiotraced not just anti-Ly6G, but also an isotype control IgG antibody with no avidity for any mouse epitope, which serves as an internal standard for a non-avid antibody. In **Figure 3E** (left panel), we plotted this as each organ’s amount of radioactive antibody, normalized to the antibody’s concentration in blood to eliminate the contribution of circulating neutrophils. In **Figure 3E’s** right panel (derived from the left panel), we show that the ratio of anti-Ly6G to control-IgG is 2.6 for the lungs and 0.62 for the liver, showing that marginated neutrophil concentration is ~4-fold higher in the lungs than in the liver; the spleen is intermediate between lungs and liver in this metric of marginated neutrophils.

Next, we used radiotracing to test if IV-injected nanoparticles can cross the alveolar capillary walls into the alveolar airspace (**Figure 3F**). This leverages a unique feature of the lungs, which is that the alveolar capillaries are separated by 1 micron from the alveolar airspace, and the alveolar airspace can be sampled by bronchoalveolar lavage (BAL). If nanoparticles leak between capillary endothelial cells at a high rate, they should be detected by BAL. We injected radiolabeled anti-PECAM liposomes, sacrificed 30 minutes later, and performed a BAL and blood draw (from the IVC). We found that there was almost zero radioactivity in the BAL, suggesting the nanoparticles do not leak out of the capillaries more than 1 micron (**Figure 3F**). In contrast, the lungs had a very high radioactivity concentration, and the blood had ~60x lower radioactivity concentration.

Last, we used flow cytometry to quantify nanoparticle uptake in *marginated* neutrophils compared to uptake in other neutrophil and leukocyte populations. As diagrammed in Figure 3F, we IV-injected fluorescent anti-PECAM liposomes, 30 minutes later injected fluorescent anti-Ly6G to label *intravascular* neutrophils, and sacrificed 5 minutes later so that the anti-Ly6G did not have time to extravasate, perfused the lungs to remove *circulating* neutrophils, and finally performed a BAL to sample the airspace as well. These techniques have been validated by other groups as allowing the isolation of each neutrophil compartment ^69,70^. We then disaggregated the lungs into a single-cell suspension and ran flow cytometry. Among all collected neutrophils that had taken up the fluorescent nanoparticles (**Figure 3G**), <u>84% were</u> <u>marginated neutrophils, 15% were interstitial neutrophils</u> (residing between the capillary and alveolar epithelium)(**Figure 3H**), and <u><1% were airspace neutrophils</u> (residing on the alveolar epithelial surface). Further, the amount of nanoparticles taken up per cell (judged by the mean fluorescence intensity [<u>MFI]) was highest among marginated neutrophils</u>, and indeed much higher than the circulating neutrophils in whole blood (**Figure 3I**).

### Mechanisms of marginated neutrophil uptake of “endothelial-targeted” nanoparticles

To elucidate the mechanisms by which marginated neutrophils take up these nanoparticles, we performed kinetic studies and engineered variations of the nanoparticles. We examined the expression of targeting epitopes on various lung cell types. First, as shown in **Figure 4A**, anti-ICAM liposome uptake in the lungs is rapid (less than 10 minutes), and only slightly decreases over one hour. The endothelial-to-neutrophil ratio remains nearly constant (**Figure 4B**). Thus, neutrophil uptake of “endothelial-targeted” liposomes is rapid and equally fast as endothelial uptake, and it is not due to the late transfer of nanoparticles between cells.

**Figure 4:**
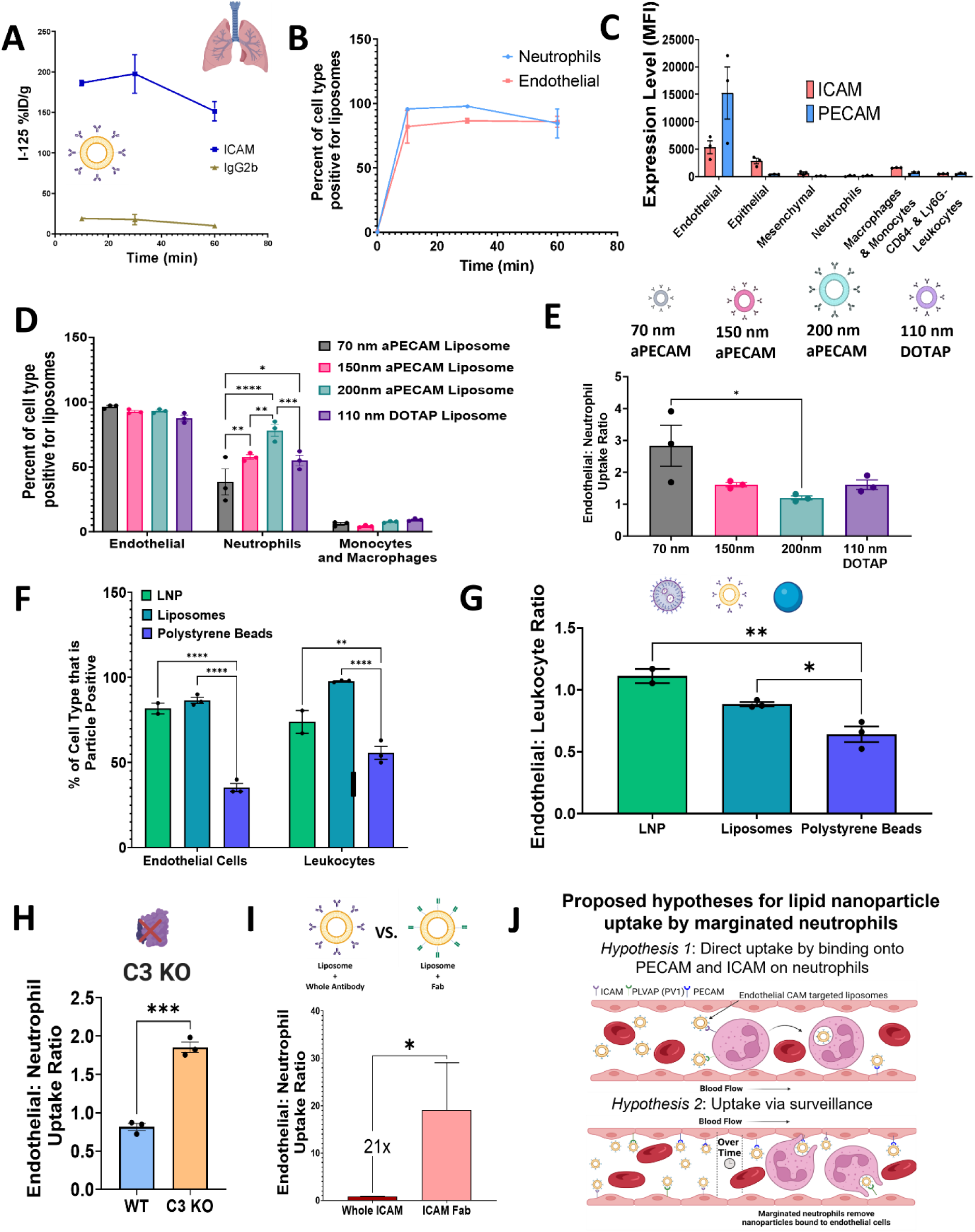
Mechanisms underlying neutrophil uptake of lung-targeted nanoparticles. **A.** Kinetics of lung uptake of radiolabeled anti-ICAM liposomes, showing it is fast (< 10 minutes), with a slight decline by 1 hour. **B**. The ratio of liposome-positive neutrophils to liposome-positive endothelial cells also declines slightly over 1 hour. **C**. Expression of ICAM and PECAM proteins on various cells in the lungs (which had not received nanoparticles), measured by flow cytometry and presented as median fluorescence intensity (MFI) **D.** In the lungs of mice that received IV-injected nanocarriers targeted to the endothelium, neutrophil uptake of CAM-targeted liposomes increases with nanoparticle size. Further, cationic liposomes with known tropism for pulmonary endothelium have similar uptake into marginated neutrophils as do CAM-antibody-targeted liposomes. **E.** Endothelial-to-neutrophil uptake ratio (derived from the experiment in D), showing larger-sized particles (200 nm) and cationic liposomes experience nearly co-equal uptake by neutrophils and endothelial cells. **F.** Neutrophil uptake of anti-ICAM nanoparticles is not specific to liposomes, but generalizes to multiple particle types, here RNA-containing lipid nanoparticles (LNPs) and polystyrene nanoparticles. **G.** The endothelial-to-neutrophil nanoparticle uptake ratio does differ somewhat between particle types, but in no case is much above 1 (meaning endothelial cells never have significantly better uptake than neutrophils). **H**. Next, we measured the effect complement opsonization of nanoparticles has on neutrophil uptake of aPECAM liposomes. We injected 10 mg/kg AF594-labeled aPECAM liposomes into complement-deficient (C3 knockout) mice and wild-type control. We observe that knocking out complement increases the endothelial-to-neutrophil uptake ratio by >2-fold. **I.** Next, we tested whether the Fc region’s presentation on a liposome’s surface affected neutrophil recognition. To test this, we generated liposomes covalently conjugated with whole anti-ICAM antibody (YN1) or antibody fragments ICAM F(ab) generated from YN1. Removal of the Fc region increases the endothelial-to-neutrophil uptake ratio by 21x. This suggests that an Fc on nanoparticles is insufficient for neutrophil uptake (IgG2b-liposomes are not taken up). Still, when present with an antibody avid for a pulmonary epitope, the Fc promotes neutrophil uptake. **J.** Updating our schematic based on our findings, we propose two hypotheses for neutrophil uptake of endothelial CAM-targeted liposomes. First, direct binding of CAM-targeted liposomes to neutrophils, and second, after liposomes successfully associate with endothelial surfaces, marginated neutrophils in their surveillance role come and remove nanoparticles and prevent uptake by endothelial cells. **Statistics**. n=3 per group unless otherwise stated. Figs. 4D, F and H, 2-way ANOVA with Tukey post hoc test *p<0.05, **p<0.01,***p<0.001, ****p<0.0001. Fig 4E,F One-way ANOVA with Tukey post hoc test *p<0.05, **p<0.01, (Fig 4H and I), Students’ T-test, *p<0.05, ***p<.001.

Next, we tested whether the targeting antibody binding to the target epitope (here ICAM and PECAM) was present on endothelial cells but not on neutrophils. We measured ICAM and PECAM levels in lung cells to evaluate this possibility using fluorescent anti-ICAM and anti-PECAM antibodies. As shown in **Figure 4C**, ICAM and PECAM levels on the endothelial cells are several orders of magnitude higher than on neutrophils.

Next, we tested the effects of size and charge on nanoparticle cell uptake into marginated neutrophils. We tested this directly by generating anti-PECAM (aPECAM) liposomes in three different sizes: 70 nm, 150 nm, and 200nm (**Figures 4D and E**). This was achieved by developing these liposomes on a Nanoassembler from Precision Nanotechnology. This allowed us to directly compare the same formulation while adjusting the flow rate to achieve differently sized nanoparticles. We discovered that larger nanoparticles have greater uptake in marginated neutrophils in the lungs, while size has little effect on endothelial uptake (**Figure 4D**). The result was that a ~3x increase in nanoparticle diameter led to a ~2x decrease in the endothelial-to-neutrophil uptake ratio (**Figures 4D-E**). This fits with prior literature showing that *macrophages* generally take up larger particles more efficiently in this size range. Still, knowing that the same holds for marginated neutrophils is very useful.

To test the effect of zeta potential on marginated neutrophil uptake, we generated a liposome containing the permanently cationic lipid DOTAP. As it is widely known that cationic nanoparticles have natural tropism to the lung endothelium ^71^, this formulation did not receive anti-PECAM targeting. Thus, these positively charged liposomes allowed us to investigate another mechanism of endothelial targeting, besides simply anti-endothelial antibodies, to determine if marginated neutrophil uptake of endothelial-binding nanoparticles is a very general phenomenon. Interestingly, we found that DOTAP-targeted liposomes have approximately the same endothelial-to-neutrophil uptake ratio as anti-PECAM-targeted liposomes. This shows that the marginated neutrophils take up “endothelial-targeted” nanoparticles, regardless of the mechanism of targeting (**Figure 4E**).

Next, we investigated whether this effect is specific to liposomes, or occurs with other nanomaterials (**Figure 4F**). For all nanomaterials tested, the same antibodies were conjugated via SATA-maleimide chemistry to achieve 50 mAb/particle. Conjugation efficiency was determined as described in the methods. First, we conjugated anti-ICAM antibodies onto RNA-lipid nanoparticles (LNPs), similar in formulation to the COVID-19 mRNA vaccines. The endothelial-to-leukocyte ratio (**Figure 4G**) was similar between RNA-LNPs and liposomes. To check if this was an effect of lipid-based nanoparticles, we next performed the same experiment, but with anti-ICAM conjugated polystyrene beads using the same conjugation chemistry, and once again observed uptake into marginated leukocytes. In our experiments, the near-equal degree of proportional uptake between endothelial cells and neutrophils was a general feature of nanoparticles targeted to endothelial-expressed epitopes, largely independent of nanomaterial.

Next, we tested the effect of serum opsonins on our endothelial-targeted liposomes, as opsonins and other serum proteins can significantly alter nanoparticle fate^71–74^. Notably, we compare all liposome formulations to IgG control liposomes, as such they should have similar protein coronas. To assess the effect of opsonins on our liposomes, we assessed the most important class of opsonin for determining nanoparticle distribution: complement proteins. Previously we have shown that complement proteins can substantially impact the biodistribution of targeted nanoparticles.^41,75–77^

We began with *in vitro* studies to determine if complement proteins were indeed opsonizing our nanoparticles. We used a nanoparticle tracking analysis (NTA) assay (**Supp Videos 6-8)** that we previously validated for measuring the opsonization of nanoparticles by the main complement opsonin, C3^41,77^. After serum incubation, IgG, aPECAM, and aICAM liposomes all modestly increased in size but were not significantly different from one another, showing corona formation (**Supp Fig 15B and C**). We further assessed the fraction of total C3-positive particles in serum. Interestingly, we found that both aPECAM and aICAM liposomes had a higher fraction of C3-positive particles, at 3% and 5% of liposomes being C3-opsonized, compared to IgG control at 1% (**Supp Fig 15E**), meaning that aPECAM and aICAM liposomes are more likely to stimulate the complement cascade.

Seeing this, we next tested if our liposomes did indeed activate the complement cascade. To do this we assessed levels of C3a in serum via ELISA assay (**Supp Fig 15F**). C3a is the byproduct of C3 catalysis and is implicated in complement-associated pseudoallergy (CARPA). We found that bare (no antibody on the surface, but still PEGylated), IgG, and aPECAM liposomes did not show significant elevation of C3a concentration compared to untreated control serum. However, we did find that aICAM liposomes had a C3a concentration above naive serum, indicating that it may cause some level of C3-mediated inflammatory response.

We next assessed the effect of complement on leukocyte uptake of aPECAM liposomes in the lung as we had established that it did not induce a CARPA-type phenotype *in vivo* and showed co-equal uptake of nanoparticles by neutrophils. To do this we injected fluorescently labeled aPECAM liposomes and compared endothelial and neutrophil uptake via endothelial-to-neutrophil ratio in both wild-type (C57/B6 mice) and C3 knockout (KO) mice (derived from C57/BL6) (**Figure 4H**). We found that C3 KO mice had a 50% reduction in neutrophil uptake of aPECAM liposomes compared to control mice, suggesting that complement opsonization is a major stimulus for neutrophil uptake of ‘endothelial-targeted’ nanoparticles.

Finally, we investigated whether there are components of the nanoparticle targeting moieties necessary for neutrophil uptake. In particular, we wondered whether the Fc region (fragment crystallizable domain) of the targeting antibody (here, anti-ICAM) is necessary for neutrophil uptake. To test this, we compared liposomes conjugated to either whole anti-ICAM monoclonal antibody, or anti-ICAM fragment antigen binding region (Fab) (**Figure 4I**). We found that Fab anti-ICAM had a 21-fold increase in the endothelial-to-neutrophil uptake ratio, but does not fully ablate it (**Supp Fig 18**). This clearly shows that neutrophil uptake does not require an Fc, but is greatly augmented by the presence of Fc’s.

This leaves two hypotheses about how marginated neutrophils in the lung take up endothelial-targeted nanoparticles (**Figure 4J**). The first hypothesis: Direct uptake by binding onto PECAM and ICAM on neutrophils. In **Figure 4C**, we found that ICAM and PECAM levels are present on endothelial and neutrophil surfaces. However, this is likely not the only mechanism responsible for this uptake. ICAM and PECAM presentation on endothelial cells is several orders of magnitude higher than on neutrophils. This gives rise to the second hypothesis: Uptake via surveillance by neutrophils of the endothelial surface. Alongside the results of **Figure 4D**, one could hypothesize that endothelial-targeted nanoparticles bind to endothelial cells. The marginated neutrophils phagocytose the nanoparticle from the endothelial surface, aided by their recognition of complement opsonins and Fc portions of the targeting antibody as depicted in **Figure 4J**.

### Marginated neutrophils take up ‘pulmonary endothelial targeted’ liposomes

To better understand the mechanism by which neutrophils take up our endothelial-targeted liposomes, we needed to visualize neutrophils’ uptake of nanoparticles in real-time. To do this, we performed *in vivo* microscopy in breathing mice to visualize nanoparticle uptake over time (**Supp Videos 2-4**). Briefly, mice were placed on a ventilator, followed by a thoracotomy, where a confocal video microscope was positioned on the surface of the lung, immobilized on the coverslip of a thoracic window by vacuum. Mice were injected with fluorescent PECAM and Ly6G antibodies to label endothelial cells and neutrophils. The videos begin by observing marginated neutrophils in the alveolar capillaries 2 minutes before the injection of nanoparticles. We then continuously record for 30 minutes after the injection of nanoparticles. As they are injected, we can observe that liposomes first deposit on the capillary surface, as shown by the representative stills in **Figure 5A**.

**Figure 5:**
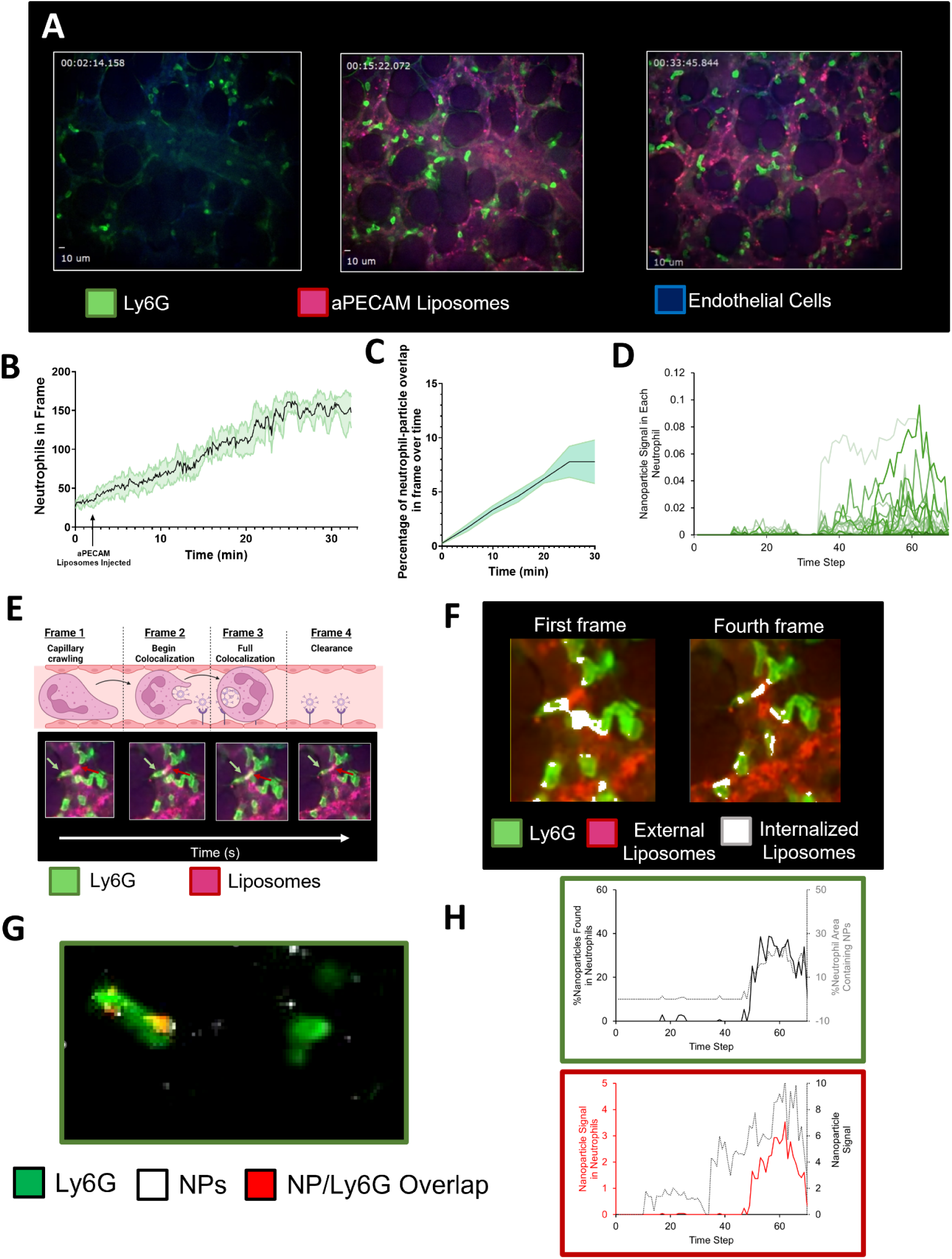
Real-time imaging in breathing mice of marginated neutrophils taking up ‘pulmonary endothelial-targeted’ liposomes. **A.** Representative still image from intravital microscopy videos shown in **Supplemental Videos 2-4**. **B.** Quantification of neutrophil movement into the frame of the video taken over time shows a steady increase in neutrophil accumulation over time after injection of aPECAM liposomes, with a reduction in the arrival rate after 20 minutes. **C.** The percentage of neutrophil-particle overlap in-frame over time, shows a gradual increase over time, as neutrophils begin to take up aPECAM liposomes in the lung. **D.** Quantification of the nanoparticle signal in neutrophils over time shows that there is a delay from when nanoparticles land on an endothelial surface and engulfment by marginated neutrophils. **E.** Representative still frames of neutrophils taking up aPECAM liposomes depicts deposition of liposomes on the endothelial surface (Frame 1), migration of neutrophils towards the nanoparticles (Frame 2), engulfment by neutrophils (Frame 3), and clearance (Frame 4). **F.** Comparing the first and fourth from **E**, investigating liposomes taken up by marginated neutrophils (liposomes in white, neutrophils in green) versus liposomes on the endothelial surface (in red) shows the active internalization of liposomes by marginated neutrophils. **G.** A representative still from **Supp Video 5.** Here liposomes on the endothelial surface colored in white, liposomes internalized neutrophils (neutrophils colored green, internalized liposomes in red) showing the direct uptake of liposomes from endothelial cells by neutrophils. **H.** Quantitative analysis of **G** assessing the percentage of nanoparticles found in neutrophils overlaid with the percentage of neutrophil area containing nanoparticles (green box). Shows that over the time step of the recording there is a delay before uptake of nanoparticles by marginated neutrophils. This is similarly shown below (red box), where nanoparticle signal nanoparticle signal is observed in early time steps, however the nanoparticle signal within neutrophils occurs only at later time steps, suggesting endothelial deposition.

Through these videos, we can very clearly observe marginated neutrophils taking up ‘endothelial-targeted’ nanoparticles. We can also notice concurrent changes in the neutrophils. First, the number of marginated neutrophils increases in the capillaries (**Figure 5B**). This fits with the long-established notion that marginated neutrophils are a “dynamic pool,” meaning that there are constant fluxes of neutrophils into and out of the marginated space, with an equilibrium number that can increase upon detection of pathogens. We further observe changes in the crawl speed (**Supp Fig 17C**), and the marginated neutrophils taking up the nanoparticles (**Figure 5C**).Quantitative analysis of these neutrophils over time (**Fig 5D**) shows a delay in uptake by neutrophils after deposition on capillary endothelial cells. This suggests deposition of nanoparticles on the endothelial surface followed by neutrophil engulfment. Indeed, **Figure 5E and Supp Fig 17A** provide multiple observations supporting this. Across four panels, we can observe a neutrophil, highlighted by green arrows, beginning to engulf deposits of aPECAM liposomes, marked in red arrows, on the capillary lumenal surface. Additional analysis of the first and fourth frames from **Figure 5E** were performed (**Figure 5F**), done by identifying the liposomes inside neutrophils (colored white) and those still on the endothelial surface (colored red). These images show neutrophils engaging in uptake of liposomes already have taken up nanoparticles, suggesting additional mechanisms are active, potentially direct binding to available cell surface markers. However, additional studies will need to be done to fully elucidate all possible mechanisms of neutrophil uptake of nanoparticles within capillaries. We further isolated an instance of neutrophil uptake from endothelial cells (**Supp Video 5 and Fig 5G**). Quantitative analysis of this video (**Figure 5H**) shows that even though liposomes signal (red box) begins early in the time step, neutrophil uptake and engulfment does not begin until the end of the time step, suggesting uptake from deposition of liposomes on the endothelial surface.

### Human lungs also display strong marginated leukocyte uptake of “endothelial-targeted” nanoparticles

To determine if the above findings generalize to humans, we tested human anti-ICAM conjugated liposomes on fresh, perfused, *ex vivo* human lungs. We regularly obtain human lungs that come from organ donors whose lungs were rejected for transplantation. As we have published multiple times before, these lungs are used within 12 hours of explantation (the same as for actual lung transplant) and are inflated, connected to an oxygenator source, endovascularly cannulated in branches of the pulmonary artery, and then perfused with a similar perfusate as used in many lung transplants ^41,42^ (**Figure 6A**). Here, we injected anti-human-ICAM liposomes into the cannulated pulmonary artery branch, perfused the lung, and created a single-cell suspension for flow cytometry. As shown in Figure 6B, marginated neutrophils had strong uptake, but less than endothelial uptake in this case. Also, slightly different than *in vivo* mice, macrophage/monocyte uptake was higher than neutrophils. The total liposome-positive cell counts and MFI were similar between neutrophils and endothelial cells and non-neutrophil leukocytes (**Figure 6B**). These results show that marginated leukocytes strongly take up “endothelial-targeted” targeted nanoparticles across species.

**Figure 6.**
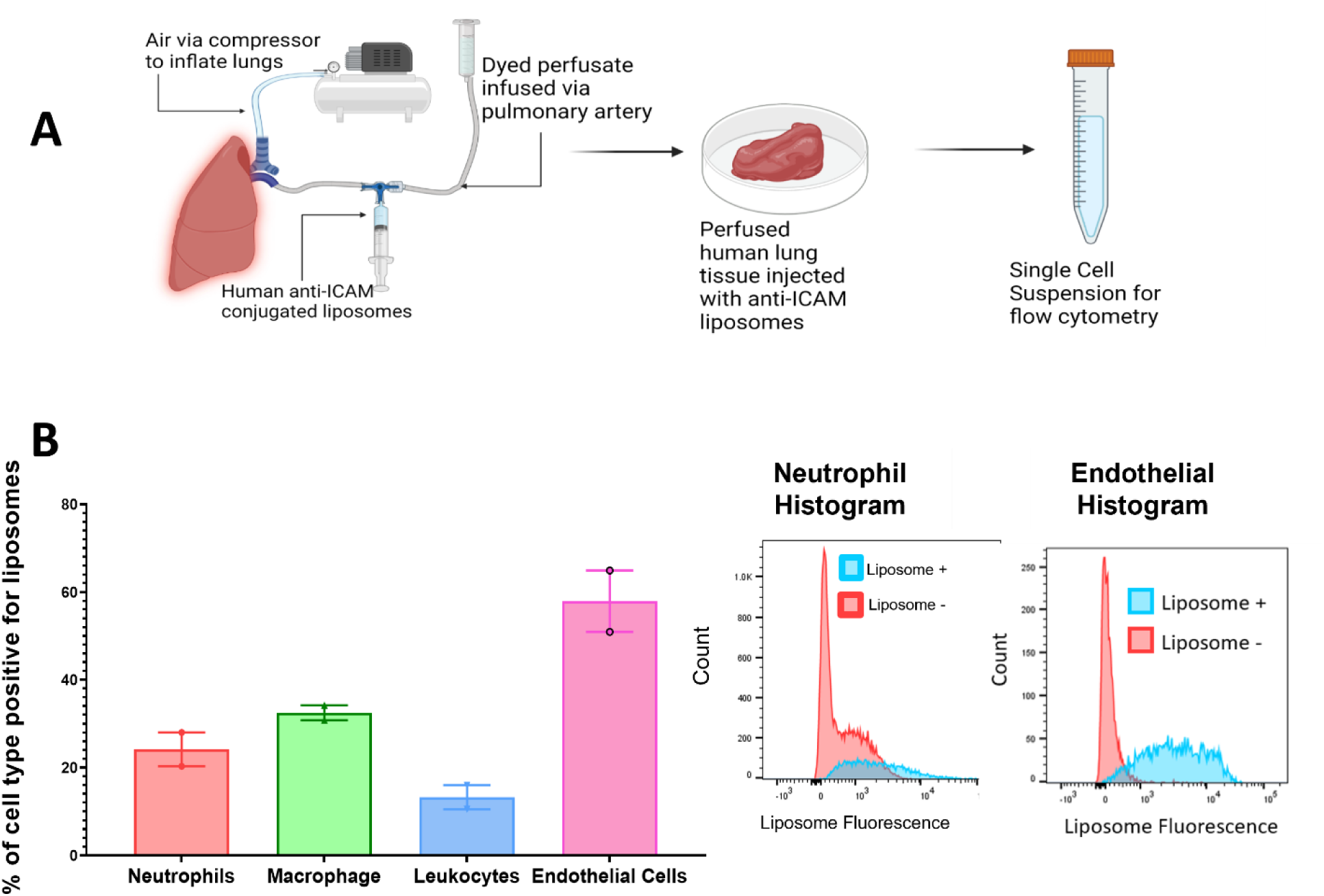
Human lungs ex vivo also display neutrophil uptake of targeted liposomes. **A.** Human lungs rejected for transplantation were oxygenated and endovascularly cannulated via the pulmonary artery for perfusion. Anti-ICAM conjugated liposomes were infused endovascularly, followed by washout with Steen’s solution and infusion of tissue dye to identify the perfused lung region for harvest. The perfused lung tissue was collected and processed into a single-cell suspension for flow cytometry. **B.** As in mice, anti-ICAM liposomes were taken up by endothelial cells and neutrophils. However, the endothelial-to-neutrophil uptake ratio was higher in humans than in mice. Histograms for neutrophils and endothelial cells are compared to liposome-negative samples, i.e., human lung samples that did not receive liposomes from the same donor.

### Inflammation changes how marginated neutrophils distribute between organs, with the lungs’ marginated neutrophils still playing a dominant role in nanoparticle uptake

Having shown that marginated neutrophils in the lungs take up endothelial-targeted nanocarriers in healthy lungs, we next wanted to see if the same held true during inflammation. It is known that marginated neutrophils increase their dwell time and numbers in the lungs during acute lung inflammation and other inflammatory diseases ^54,55,57^. However, it is not known how such pathologies affect nanocarrier uptake.

To investigate this, we first started by creating an inflammatory injury *remote* from the lungs, with cecal ligation and puncture (CLP; which is also a model of sepsis). Two hours after injury, we IV-injected radiolabeled anti-Ly6G antibodies (specific to neutrophils), allowed them to circulate for 5 minutes, then measured radioactivity in each organ (the short time prevents antibody extravasation into parenchyma). In **Figure 7A**, we plot each organ’s concentration of radioactivity, normalized to the concentration in blood to eliminate the contribution of circulating neutrophils. Figure 5A shows that the lungs have a higher concentration of marginated neutrophils at baseline than the liver, and the lung concentration of marginated neutrophils increases dramatically upon systemic inflammation. Next, we measured the uptake of “endothelial-targeted nanoparticles” in these marginated neutrophils. We injected fluorescently labeled anti-PECAM liposomes, then 30 minutes later IV-injected fluorescent anti-Ly6G to label both circulating and marginated neutrophils, and sacrificed 5 minutes later, followed by harvest of circulating blood, and perfusion to remove circulating neutrophils. After making a single-cell suspension of lungs and spleen, we stained the cells with a different color of anti-Ly6G to label interstitial/intraparenchymal neutrophils. In **Figure 7B** top panel, we display data from the lungs of CLP mice, showing that among liposome-positive leukocytes, by far the most abundant are marginated neutrophils. Next, we measured what fraction of neutrophils were liposome-positive in the lungs, blood, and spleen. This shows that neutrophils in the lungs and spleen were equal to each other in liposome uptake, and this liposome uptake was ~2x higher than that of circulating neutrophils in the blood sample (**Figure 7B, bottom panel**). Further, the histograms of liposome fluorescence (third panel) show that while marginated neutrophils of the lungs do not have higher liposome uptake than other neutrophil populations, they are much more abundant (on a per-tissue-volume basis). Finally, we used confocal microscopy to assess if marginated neutrophils in multiple organs are taking up nanoparticles (**Figure 7C**). These confocal images were taken after IV injection of fluorescent nanoparticles and IV injection of fluorescent anti-Ly6G shortly before sacrifice, thus labeling only marginated / blood-exposed neutrophils’ uptake of nanoparticles (**Figure 7C**). All this data confirms that: marginated neutrophils in the the lungs and spleen take up nanoparticles at higher rates than circulating (blood) neutrophils, but the very high abundance of marginated neutrophils in the lungs is the major reason the lungs take up endothelial-targeted nanoparticles much more.

**Figure 7:**
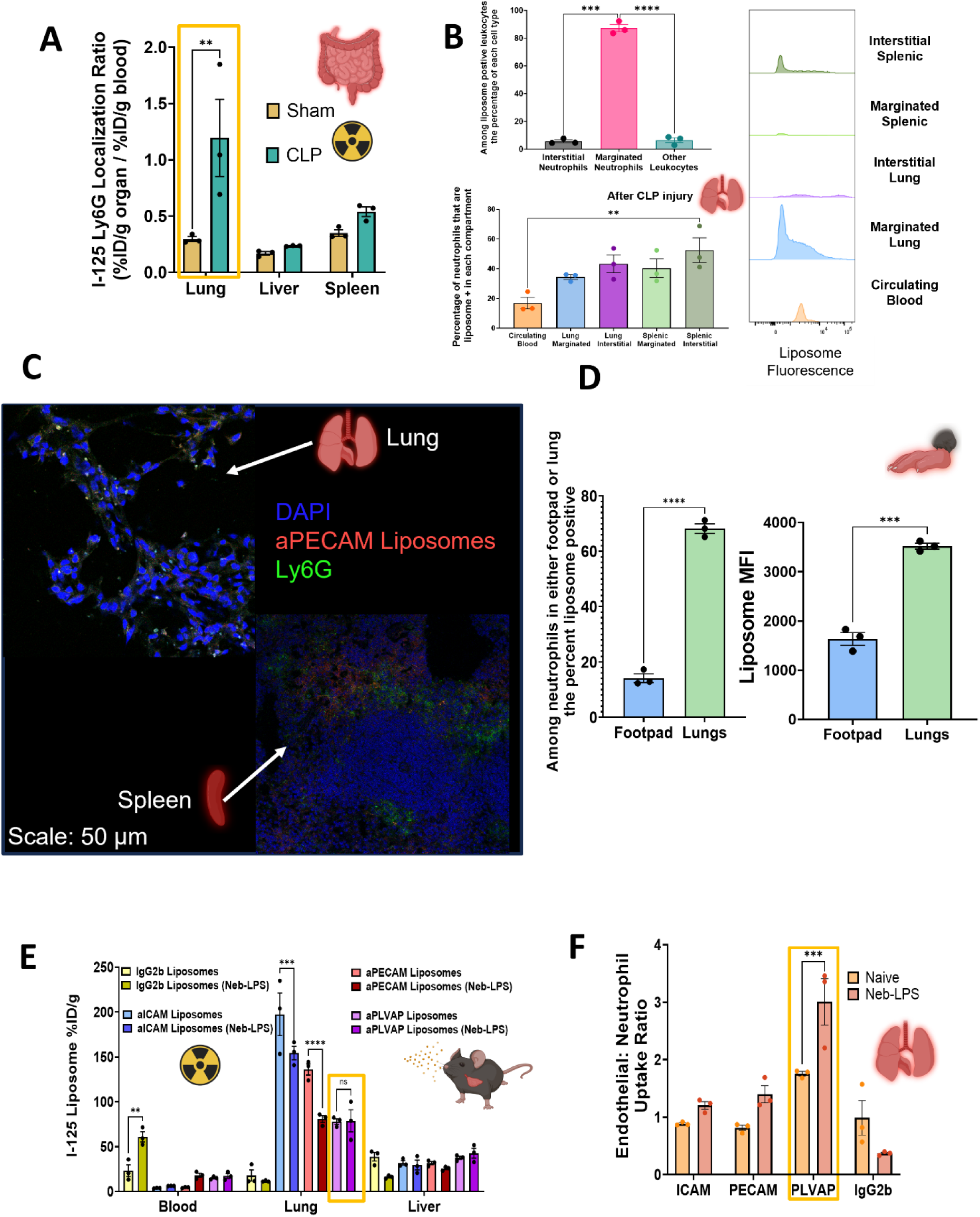
During both local and remote inflammation, marginated neutrophils in the lungs play a dominant role in uptake of nanoparticles that bind to the vasculature. **A.** Biodistribution of IV-injected I-125 radiolabeled anti-Ly6G antibody shows that 2 hours after CLP injury, neutrophil presence in the lung increases significantly **B.** CLP mice were IV-injected with fluorescent anti-PECAM liposomes and anti-Ly6G antibody (to label marginated neutrophils) via the same protocol as in Figure 3. This showed that marginated lung neutrophils take up nanoparticles to a greater extent than neutrophils in other lung compartments, blood, or spleen. **C.** Microscopy shows the marginated pool of neutrophils in the lungs colocalize well with aPECAM liposomes after CLP injury. However, in the spleen, while there is a presence of aPECAM liposome and neutrophils, there is less colocalization. **D.** In the CFA footpad inflammation model, flow cytometry reveals that ~60% of the marginated neutrophils in the lung take up fluorescently labeled aPECAM liposomes. In comparison, only ~20% of neutrophils take up nanoparticles in the injured footpad. Liposome MFI similarly shows that lung-marginated neutrophils more avidly take up aPECAM liposomes than neutrophils found in the footpad. **E.** Biodistribution of radiolabeled nanoparticles shows that ALI (nebulized LPS) decreases total lung uptake for ICAM- and PECAM-targeted nanoparticles, but not PLVAP- or isotype control. **F.** Gating first to identify cell types, then identified the percentage of each cell type that takes up our CAM-targeted liposomes. Taking these percentages for endothelial cells and neutrophils we get the endothelial-to-neutrophil uptake ratio and find that it is increased for all targeting moieties, but most for PLVAP, which was already the most endothelial-specific. **Statistics.** n=3 for all groups. Fig 7 A, E, and F, 2-way ANOVA with Tukey post hoc test **p<0.01,***p<0.001, ****p<0.0001. Fig 7B One-way ANOVA with Tukey post hoc test *p<0.05, **p<0.01, Fig 7D Unpaired T-test with Welch’s correction, ****p<.0001, ***p<0.001.

Next, we wanted to investigate if the marginated neutrophils of the lungs take up more endothelial-targeted nanoparticles than the local neutrophils in an inflamed remote organ. This was done using Complete Freund’s Adjuvant or CFA-footpad model. Using this model we found severe inflammation in the foot, and the number of local neutrophils increased (**Supp Fig 10 E, F**). However, when CFA-footpad mice were IV-injected with anti-PECAM nanoparticles, we found that the lungs’ marginated neutrophils had much higher nanoparticle uptake than the neutrophils of the inflamed foot (**Figure 7D**). Thus, the pulmonary marginated neutrophils also have a special role in filtering out nanoparticles and microbes, even compared to the local neutrophils in remote inflamed tissue.

After studying the effects of remote inflammation on marginated neutrophils in the lungs, we next investigated marginated neutrophils during *local* lung inflammation. One particularly important clinical indication for which lung endothelium-targeted nanoparticles were originally developed is acute alveolar inflammation. This occurs most commonly in acute respiratory distress syndrome (ARDS), which is the main cause of death in COVID-19. ARDS and other alveolar inflammatory conditions (e.g., post-lung transplant) cause increased margination of neutrophils in the lung vasculature and infiltration of neutrophils into the alveoli ^35–38^. Given that marginated neutrophils become much more numerous and activated in ARDS and display more ICAM-1 on their surface^15,16,25^, we hypothesized that, in inflamed lungs, neutrophils would account for even more uptake of the above lung-targeted nanoparticles. An extensive study by our group using non-targeted nanoparticles of various classes showed nanoparticle tropism to marginated neutrophils in the lung^41^.

To test this hypothesis, we employed the nebulized lipopolysaccharide (LPS) mouse model, which mimics many key aspects of ARDS. As shown in **Supp Fig 10B**, within 4 hours of nebulized LPS, neutrophils increase in the lungs by about 3-fold. We injected radiolabeled liposomes into these mice at 4 hours post-LPS, and measured total lung uptake. Surprisingly, we found that lung uptake of PECAM- and ICAM-targeted liposomes was lower in nebulized LPS (neb-LPS) mice than in naive mice (**Figure 7E**). Notably, this is different than our prior studies ^15,16^, in which ICAM-nanoparticle uptake was increased in inflamed lungs, mostly because those prior studies used the less ARDS-like models of IV-LPS or intra-tracheal instillation LPS, both of which do not cause as much neutrophil ingress into the alveolar airspace, and therefore leave their marginated neutrophils in the alveolar capillaries available for nanocarrier uptake. Importantly, in PECAM- and ICAM-targeted liposomes, the neutrophil uptake percentage decreased by a similar amount as to the total lung uptake (**Supp Fig 4C**). By contrast, LPS did not affect endothelial uptake. This confirms that the total mass of nanoparticles taken up by neutrophils in naïve mice must represent a large percentage of the total mass taken up by the lungs, since decreased neutrophil uptake is associated with a proportionate decrease in total lung uptake.

In both naive and neb-LPS mice, PLVAP targeting had a higher endothelial-to-neutrophil ratio than ICAM—and PECAM-targeting (Figure 7F). However, the ratio improved from around 2-to-1 in naive mice to 3-to-1 following nebulized LPS. PLVAP is superior to the more commonly targeted epitopes of PECAM and ICAM for the most significant endothelial-targeting specificity.

As in prior sections, we reanalyzed the data again, gating first for liposome-positive cells and then analyzing the fraction of liposome-positive cells represented by each cell type, as shown in Supp Fig 3B. For both PECAM- and PLVAP-conjugated liposomes, among liposome-positive cells, there was no change compared to naive in the fraction of neutrophils and endothelial cells (e.g., among PLVAP-liposome-positive cells, ~80% were endothelial cells in both naive and nebulized-LPS mice). However, among ICAM-liposome-positive cells, a lower percentage were marginated neutrophils, and a higher percentage were monocytes and macrophages. This is supported by the increased ICAM presentation we observed in nebulized-LPS-exposed monocytes and macrophages (Supp Fig 7A). Thus, the nebulized LPS model of ARDS somewhat changes the neutrophil uptake, but not endothelial uptake, in a manner that is specific to the targeting moiety.

## DISCUSSION

For decades, and across hundreds of papers, one of the best examples of nanomedicine targeting a single organ and single cell type was the localization of nanoparticles to the lungs using endothelial-targeted antibodies. While these were implicitly assumed to target only endothelial cells in the lungs, we have shown that their lung-targeting owes as much to neutrophil uptake as to endothelial cells. We have shown that this is true across diverse nanomaterials and targeting moieties. Indeed, we showed that marginated neutrophil uptake is not even dependent on targeting moieties at all, as nanoparticles with physicochemical tropism to the lungs provided by cationic charge are also taken up by marginated neutrophils. Further, this occurs in healthy and inflamed lungs and is augmented by remote inflammation (such as intestinal perforation) in both *in vivo* mouse models and *ex vivo* human lungs.

These results have significant implications for developing lung-targeted nanocarriers into clinically useful products. On the plus side, marginated neutrophils are widely considered bad actors in ARDS and other sterile lung inflammation, as they secrete toxic mediators (proteases, reactive oxygen species, proinflammatory cytokines, etc.) and serve as a reservoir for neutrophils to diapedese into the alveolar airspace, where they damage the lung parenchyma and impede normal respiration. Thus, targeting neutrophils with nanocarriers could be beneficial if the nanocarriers carry drugs that control the deleterious aspects of neutrophils. This has been explored in bacterial and endotoxin (LPS) models of ALI injury, taking advantage of particle properties that innately create tropism in the lung ^9,41,78–82^. By contrast, “accidental” neutrophil targeting by nanoparticles could produce major side effects. Neutrophils secrete toxic mediators in response to phagocytosis of complement and/or IgG-opsonized pathogens, and nanoparticles have material properties similar to those of these pathogens. They therefore might induce such deleterious neutrophil responses ^9,43^. Either way, knowledge of nanoparticle uptake by marginated neutrophils will be key to the translational development of nanomedicines.

To control whether lung-targeted nanoparticles produce deleterious or beneficial effects, knowing the mechanism by which such nanoparticles end up in marginated neutrophils is essential. Towards that end, our studies suggest a mechanism of neutrophil uptake of “endothelial-targeted” nanoparticles. In healthy mice, the expression of ICAM and PECAM on endothelial cells is several orders higher than on neutrophils. However, the uptake of anti-CAM nanoparticles through this direct binding of nanoparticles to ICAM or PECAM on neutrophils does not fully explain the phenomenon. Instead, we propose an additional mechanism in which anti-CAM nanoparticles first bind to either ICAM, PECAM, or PLVAP on endothelial cells and are then recognized by the marginated neutrophils as invading pathogens (in part due to Fc fragments on the nanoparticle), followed by phagocytosis the nanoparticles by the marginated neutrophils. This is strongly supported by the finding that IgG-nanoparticles are not taken up by marginated neutrophils (**Supp Fig 18**; suggesting Fc on the surface is not sufficient for uptake), and that nanoparticles conjugated to Fab-anti-ICAM have reduced neutrophil uptake. Additionally, we have shown video proof by videomicroscopy of nanoparticle ‘stripping’ from endothelial cells by marginated neutrophils (**Supp Videos 2-5**). Further, physicochemically targeting nanoparticles to the negatively charged endothelial glycocalyx produces a similar neutrophil uptake. Importantly, this mechanism fits with direct visualization of how marginated neutrophils take up pathogens in the lungs ^83^. In intravital microscopy of the lungs, IV-injected microbes are observed to deposit on the endothelium, and then within seconds to minutes, neutrophils phagocytose the microbe. This mechanism also suggests that to avoid marginated neutrophils, steps should be taken to minimize similarity to microbes, such as minimizing Fc fragment inclusion and complement opsonization (**Fig 4 H and I**).

More generally, our results suggest that true organ- and cell-type-specific targeting may not be achievable with nanoparticles. Mono-targeting of nanoparticles has long been sought, but we show here that the mono-targeting was illusory for one of nanomedicine’s best examples of this ideal. Instead, perhaps the way forward is to choose cargo drugs that themselves encode cell-type specificity, such as nucleic acids with sequences that confer cell-type-specific expression. For these reasons and more, it is our hope that the present study encourages further quantification of what cell types take up nanoparticles within a target organ, with the goal of not achieving ideal mono-targeting, but instead understanding how uptake by off-target cells can be mitigated or leveraged to benefit.

While the findings reported here have obvious implications for the future development of “endothelial-targeted” nanoparticles, they also have other important implications for nanomedicine. Most importantly, these results point out that the marginated neutrophils of the lungs might be considered as a part of reticuloendothelial system (RES) clearance of nanomedicines. The RES is classically considered to be composed, at the organ level, of only the liver and spleen, and its constituent cells are claimed to be of the monocyte/macrophage lineage (hence the alternative name for the RES: mononuclear-phagocytic system (MPS)). But we have shown here that the lungs, particularly the marginated neutrophils, play a large role in surveillance and non-specific uptake of nanoparticles, and thus should perhaps be considered a part of the RES.

How might the marginated neutrophils fit into the RES paradigm?

To explore this, it is important to define marginated neutrophils’ location within organs. Marginated neutrophils are defined as residing for prolonged times in the lumen of the microvasculature ^28,29,84^. Extensive video microscopy both in this study (**Supp Videos 2-4**) and other labs ^53,85^ have shown that while marginated in the lung capillaries, marginated neutrophils do not simply get pushed around by blood flow, but crawl around the highly anastomotic capillary network of the alveoli (it looks like a *<u>net</u>* rather than straight tubes), navigating as a car might within the grid-like roads of a city. When a pathogen lands in this net of lung capillaries, the marginated neutrophils are caught on video, quickly traversing to phagocytose the organism. This illustrates that marginated neutrophils are at least functionally distinct from neutrophils in bulk circulation. However, the marginated neutrophils have never been shown to be distinct from circulating neutrophils in terms of gene expression or surface markers. Indeed, the marginated neutrophils regularly leave the lungs into the circulation, and return, which is why they are considered a “dynamic pool.” ^55^ That being said, in this study we found that marginated neutrophils more highly expressed CD11b (**Supp Fig 16**), a marker that in neutrophils is associated with mobility and neutrophil crawling. Our data seems to suggest that these marginated neutrophils are a phenotype between interstitial and circulating neutrophils, an interesting finding that warrants further study. Notably, this dynamic nature does not change a key fact about marginated neutrophils for nanomedicine: *The marginated neutrophils’ location residing in an organ means that they can markedly change the organ distribution of nanoparticles*. For instance, we have shown here that marginated neutrophils represent a large fraction of lung nanoparticle uptake. Additionally, marginated neutrophils’ uptake of nanoparticles in the lungs may produce side effects in that organ, which may differ from the side effects produced by nanoparticle uptake into circulating neutrophils. Thus, the distinction between marginated and circulating neutrophils is a key property of nanomedicine.

The next step in understanding marginated neutrophils’ potential position in the RES is delineating how they are *distributed between organ*s. A prior basic science study imaged thick sections of many histological slides to show that the lungs have the most marginated neutrophils, followed by the spleen, liver, and skin^56^. We confirmed this with our radiotracing studies, showing marginated neutrophils were >4x more concentrated in the lungs than the liver, with the spleen in between (**Figure 3E**). Thus, marginated neutrophils are not unique to the lungs but are the most plentiful leukocyte there.

The marginated neutrophils in the lungs also take up more endothelial-targeted nanoparticles than other neutrophils. In healthy mice, the amount of nanoparticles taken up per cell (measured by MFI) was 6.5-fold higher in marginated neutrophils than the circulating neutrophils in whole blood (**Figure 3H**), showing marginated neutrophils are different than circulating neutrophils. Further, in the CFA-footpad model of skin / soft tissue inflammation, neutrophils increase in numbers in the inflamed foot. Still, these neutrophils take up endothelial-targeted nanoparticles much less avidly than the lungs (**Figure 7D**).

However, the lungs are not totally unique in having marginated neutrophils that avidly take up endothelial-targeted nanoparticles. As judged by the number of nanoparticles taken up per cell (estimated by MFI), marginated neutrophils in the lungs are equivalent to those in the spleen, at least in mice with the sepsis-like condition of CLP. However, the lung has many more marginated neutrophils per gram than the spleen (**Figure 7A**), so total lung uptake into marginated neutrophils is higher in the lungs than in the spleen.

The next point to consider for marginated neutrophils’ position in the RES is how they change in disease states. As shown in Figure 7, marginated neutrophils’ behavior varies depending on the disease state, but in all cases, they play a prominent role in nanoparticle uptake. In local lung inflammation induced by nebulized LPS (**Figure 7F**), pulmonary neutrophils take up fewer nanoparticles than healthy mice. By contrast, in remote inflammation (abdominal or foot inflammation), pulmonary marginated neutrophils take up either more (CLP model) or unchanged amounts of nanoparticles.

Finally, the last point to consider about marginated neutrophils’ position in the RES is the mechanism by which they take up nanoparticles. Our data suggests that marginated neutrophils’ main function in the RES is to take up nanoparticles that adhere to the vasculature’s luminal surface (**Figures 3, 5, 7, and Supp Videos 2-5**). Supporting this is the finding that marginated neutrophils in the lungs take up very few nanoparticles without endothelial tropism (e.g., IgG-liposomes used as controls), but avidly take up nanoparticles that bind to endothelium, across very diverse targeting methods (cationic nanoparticles and many different endothelial-binding antibodies^11,15,16,18,25,71,86^). Further, attributes that make a nanoparticle more attractive to a local phagocyte (possessing an Fc portion of an antibody, larger diameter) increase marginated neutrophil uptake of nanoparticles, but only when the particles bind to the endothelium. Indeed, it was previously found that 500 - 2000 nm polystyrene particles are taken up by neutrophils (type not studied) in the lungs; likely this uptake was marginated neutrophils phagocytosing these large particles after they got lodged in the net-like labyrinth of alveolar capillaries^79^. Finally, our finding that marginated neutrophils display their strongest nanoparticle uptake in the lungs (as opposed to the liver or spleen), fits with the special anatomic features of alveolar capillaries, which form a grid-/ / net-like pattern that allows the neutrophils to crawl retrograde to phagocytose particles bound to the endothelium.

Thus, to summarize the position of the marginated neutrophils in the RES paradigm: Marginated neutrophils are most abundant in the lungs, but also prominent in the spleen. These cells are unique from circulating neutrophils in that they can capture and hold nanoparticles in the organ where the marginated neutrophils reside. Marginated neutrophils do not take up all nanoparticles, but only ones that bind to or lodge in the vascular lumen, regardless of what causes that binding. Therefore, marginated neutrophils play a significant role in clearing the blood of the broad class of nanoparticles that bind vasculature, a key role of the RES.

## Notes

### Competing Interest Statement

The authors have declared no competing interest.

