## Supplemental Figures for "Marginated neutrophils in the lungs effectively compete for nanoparticles targeted to the endothelium, serving as a part of the reticuloendothelial system"

### Marginated neutrophils in the lungs are a major contributor to uptake for nanoparticles targeted to endothelial cells

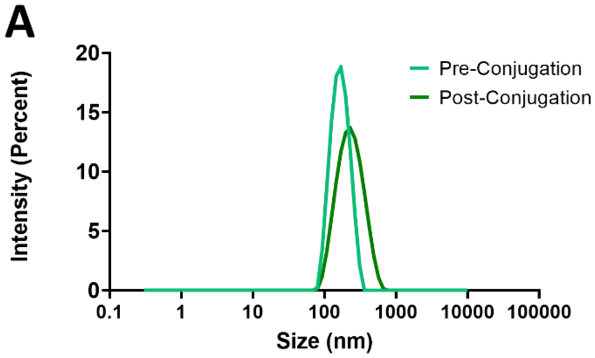


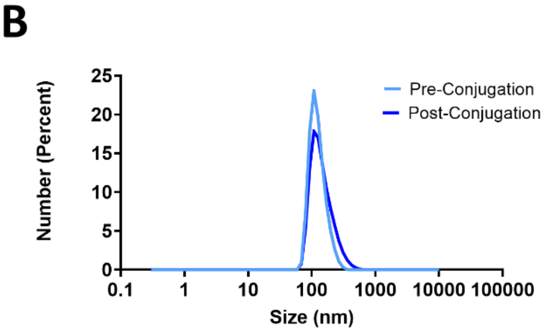


| Parameter | Pre-Conjugation | Post-Conjugation |
|---|---|---|
| Z-average (nm) | 158.5 | 207.9 |
| Polydispersity Index (PDI) | 0.06595 | 0.1448 |
| Peak mean by intensity | 171.7 | 245.4 |

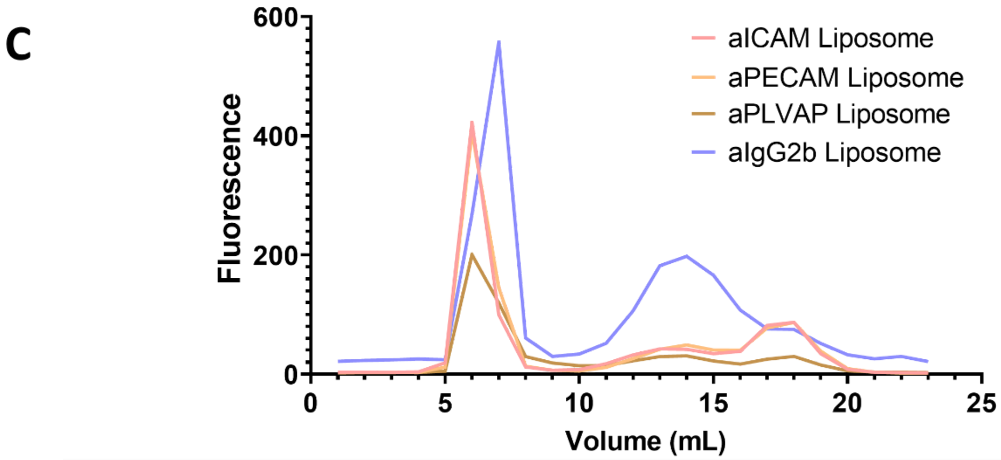


| Formulation | Conjugation Efficiency | mAb per liposome |
|---|---|---|
| **aICAM Liposome** | 79% | 49 |
| **aPECAM Liposome** | 78.1% | 49 |
| **aPLVAP Liposome** | 78% | 49 |
| **aIgG2b Liposome** | 47% | 29 |

**Supp Fig 1. Characterization of monoclonal antibody conjugated liposomes A.** Intensity peaks show that after conjugation with monoclonal antibodies, mAb-conjugated liposome size does not fluctuate significantly and does not indicate aggregation. Similarly, in **B., the number of distribution particles in DLS is** unaffected. **Supp Table 1** shows changes in nanoparticle size and PDI pre- and post-conjugation **C.** Elution profile of mAb-conjugated liposomes. Based on Sepharose size exclusion chromatography, we anticipate mAb-conjugated liposomes to elute in the first peak, with unbound and free fluorescence in subsequent peaks. This is used to generate **Supp Table 2.** A calculation of conjugation efficiency and achieved antibodies on the surface of the liposomes.

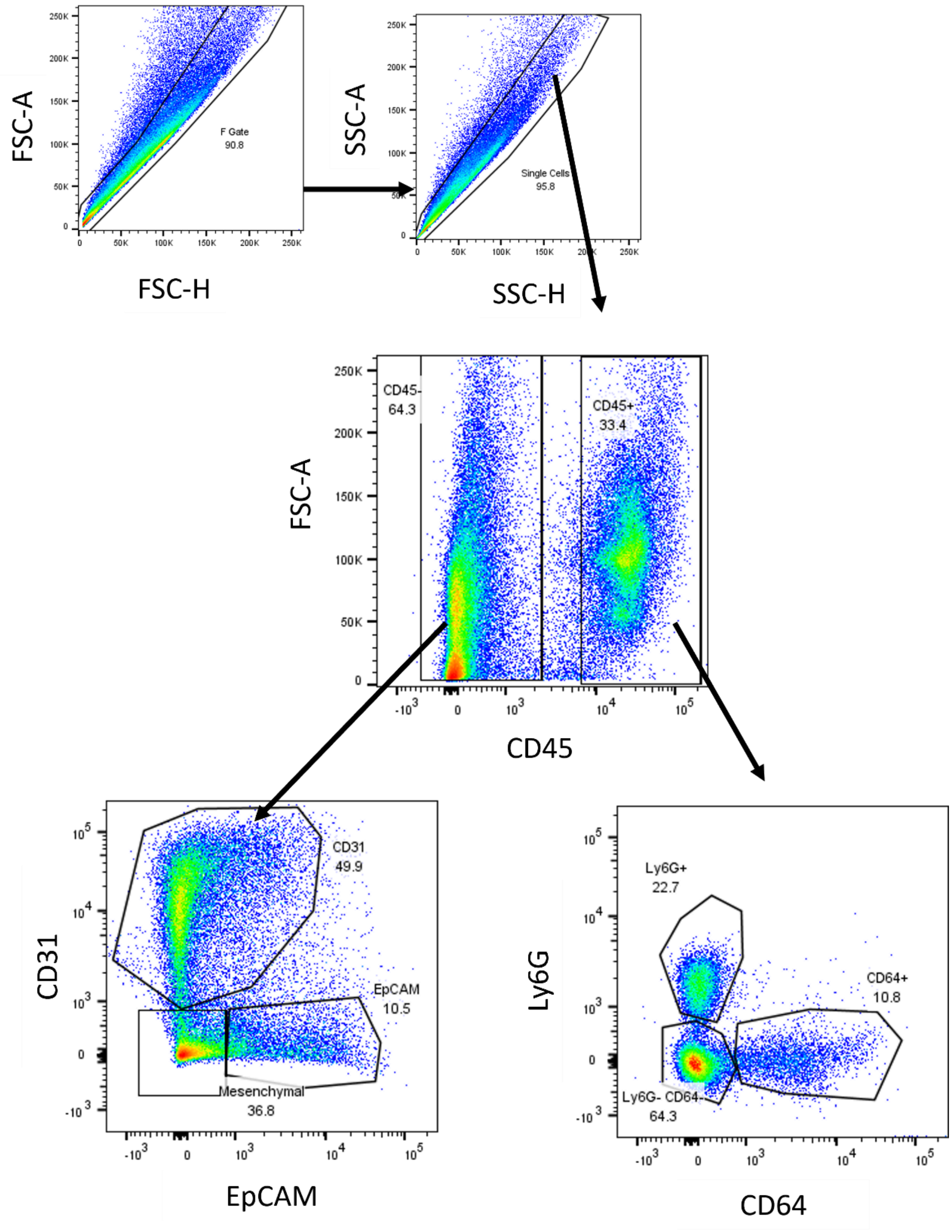


**Supp Fig 2. Gating strategy used to identify key cell populations in mouse lungs**. Cell uptake of endothelial uptake was then assessed by seeing the percentage of cells that were liposome-positive in these groups. Here we identify neutrophils (Ly6G+), monocytes and macrophages (CD64+), endothelial cells (CD31+), and epithelial cells (EpCAM+)

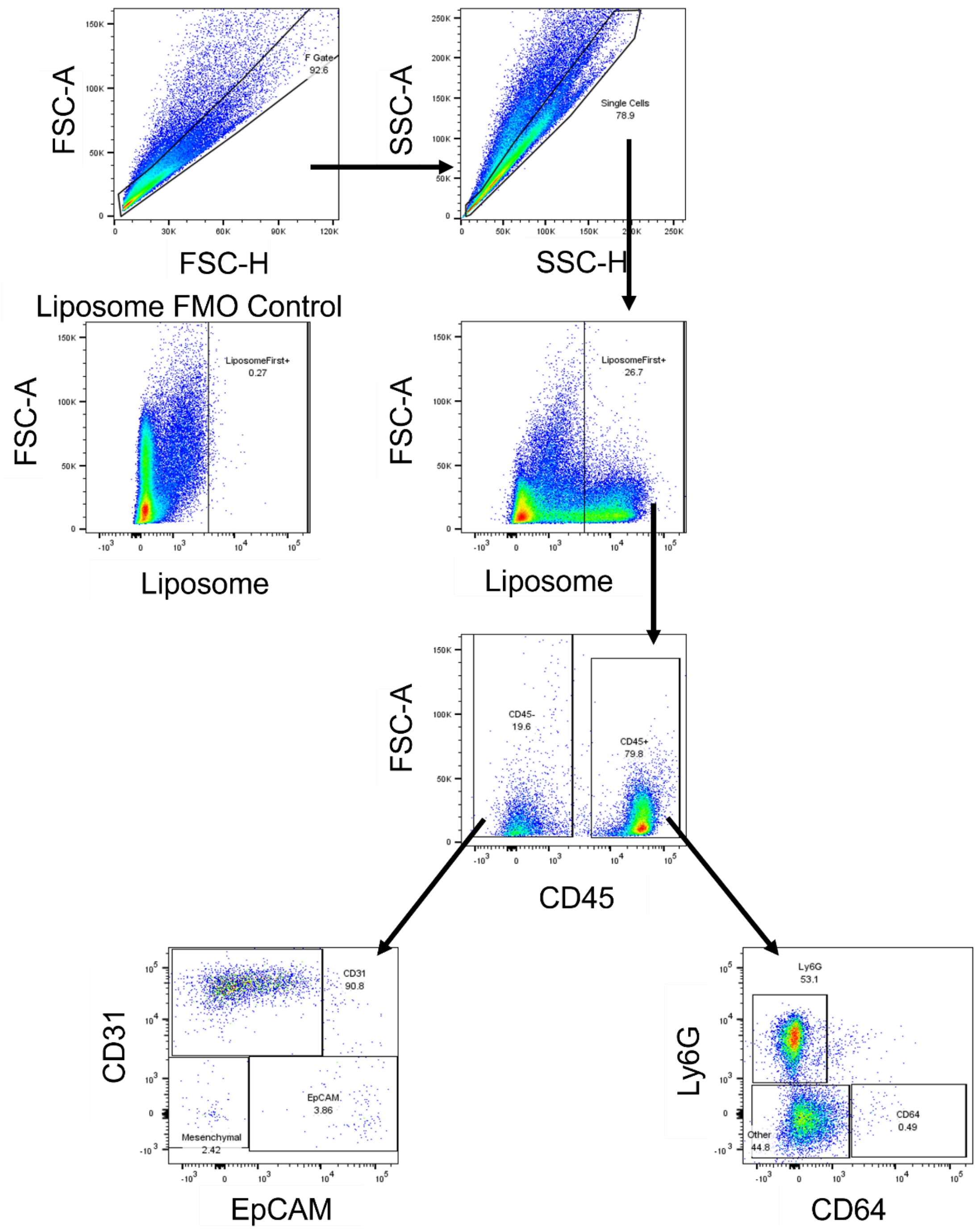


**Supp Fig 3. The gating strategy employed for identification of the fraction of liposome-positive cells**. To identify the liposome-negative population, a fluorescence minus one (FMO) control was performed by surface staining for all cell types as previously described in methods in mouse lungs that were not provided fluorescent liposomes. Here we identify neutrophils (Ly6G+), monocytes and macrophages (CD64+), endothelial cells (CD31+), and epithelial cells (EpCAM+)

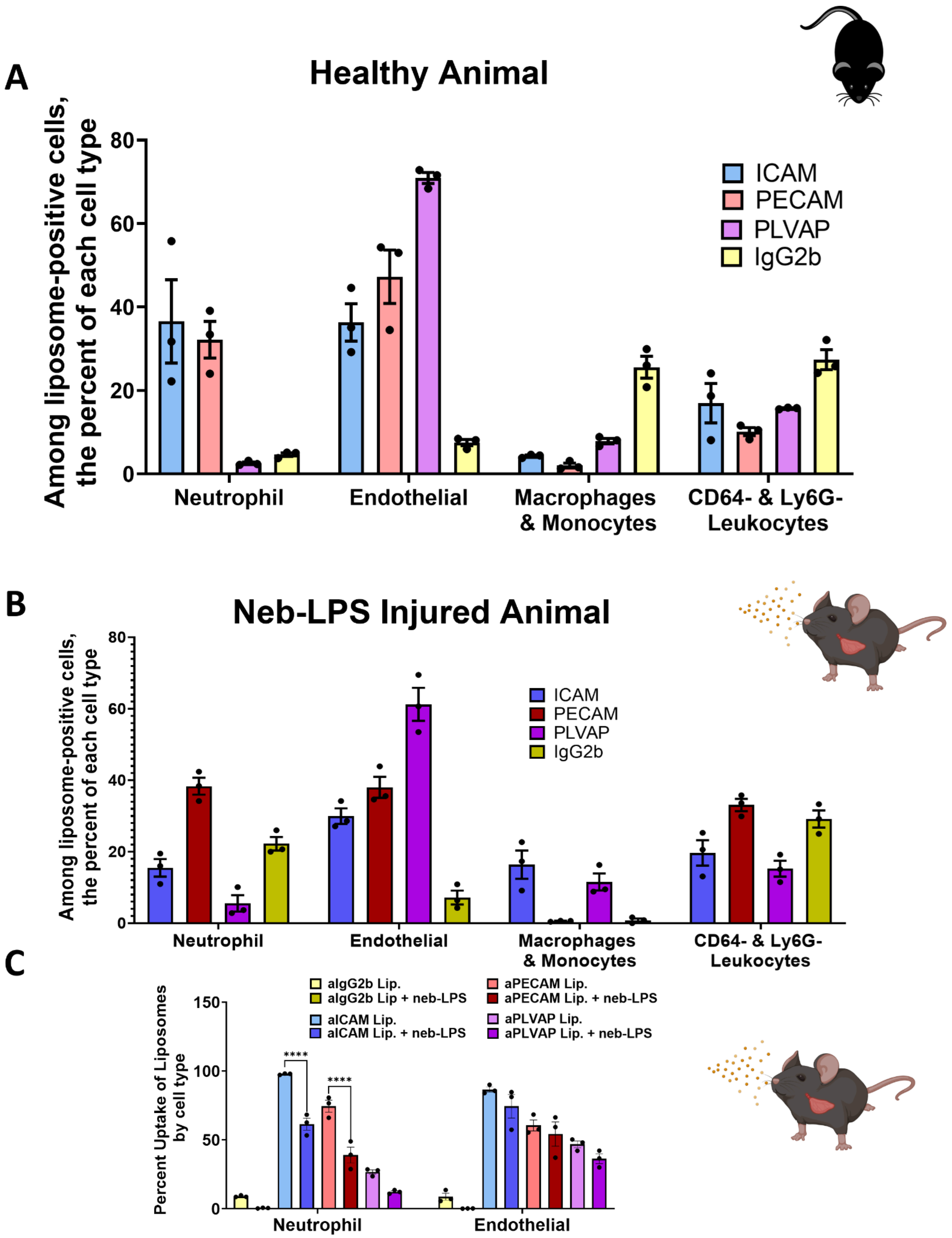
A
Healthy Animal
Among liposome-positive cells, the percent of each cell type
80
60
40
20
0
ICAM
PECAM
PLVAP
IgG2b
Neutrophil
Endothelial
Macrophages & Monocytes
CD64- & Ly6G- Leukocytes
B
Neb-LPS Injured Animal
Among liposome-positive cells, the percent of each cell type
80
60
40
20
0
ICAM
PECAM
PLVAP
IgG2b
Neutrophil
Endothelial
Macrophages & Monocytes
CD64- & Ly6G- Leukocytes
C
Percent Uptake of Liposomes by cell type
150
100
50
0
aIgG2b Lip.
aIgG2b Lip + neb-LPS
aICAM Lip.
aICAM Lip. + neb-LPS
aPECAM Lip.
aPECAM Lip. + neb-LPS
aPLVAP Lip.
aPLVAP Lip. + neb-LPS
****
****
Neutrophil
Endothelial

**Supp Fig 4. Breakdown of endothelial-targeted liposome-targeted cells. A.** Gating for the fraction of liposome-positive cells shows neutrophils and endothelial each take up 40% of aICAM- and aPECAM-conjugated liposomes. However, for aPLVAP-conjugated liposomes, 80% were taken up by endothelial cells. **B.** After neb-LPS injury, we find a slight reduction in endothelial uptake of aICAM-conjugated liposomes in favor of monocytes and macrophages. Endothelial cells remained the predominant cell type that takes up PLVAP-conjugated liposomes. **C.** Gating first among individual cell types, then assessing the percentage of each cell type positive for liposomes after neb-LPS treatment shows a decrease in neutrophils positive for liposomes.

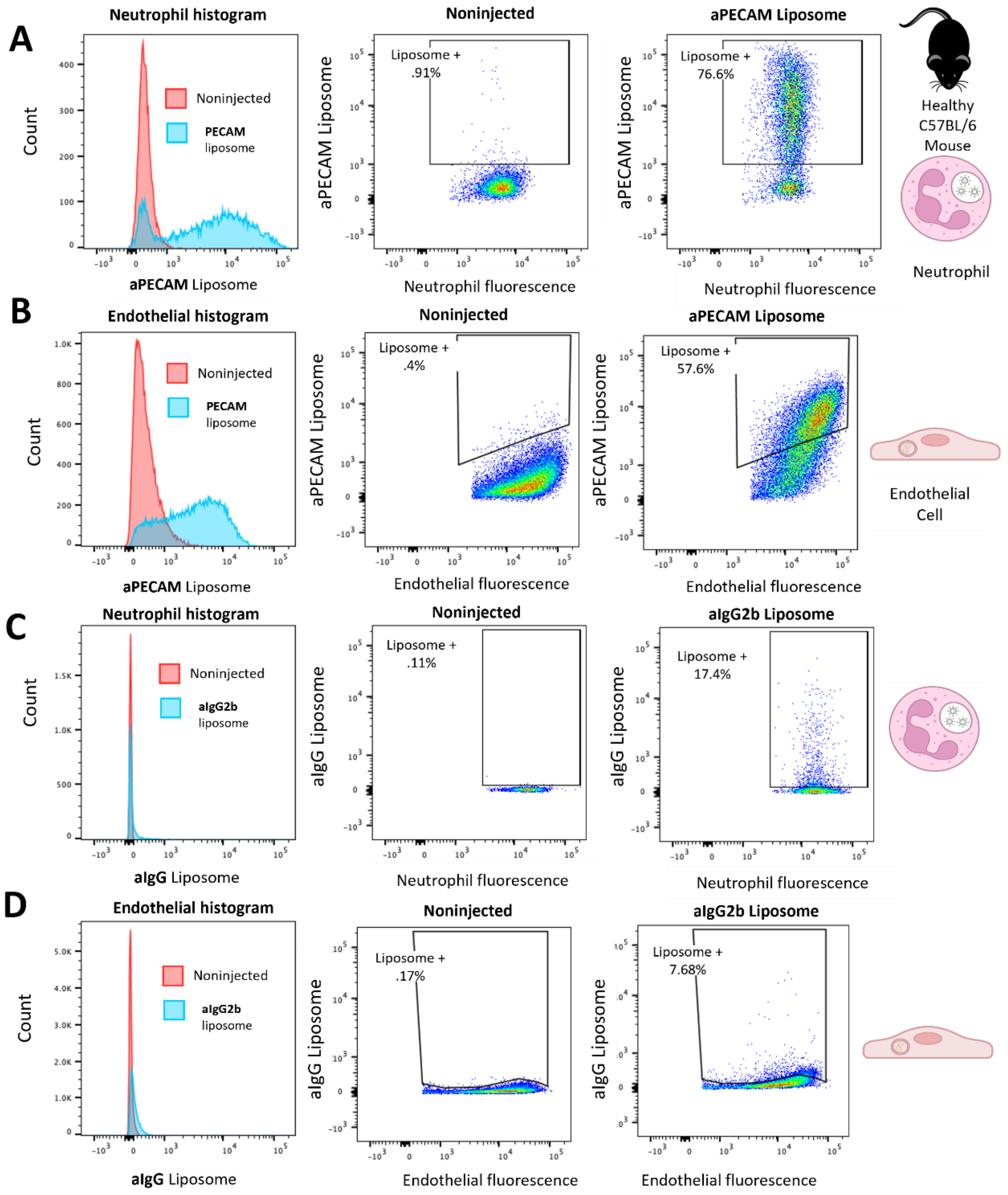


**Supp Fig 6. Histograms and representative dot plots of aPECAM and aIgG2b liposomes. A.** leftmost panel is a histogram comparing liposome uptake neutrophils in noninjected healthy animals in red to healthy animals injected with aPECAM liposomes depicted in blue. The remaining two panels are representative dot plots of noninjected control (middle panel) serving as a fluorescence minus one (FMO) control for mice injected with aPECAM liposomes. **B**. Same as panel A but represents uptake of aPECAM liposomes by endothelial cells. Similarly, **C** and **D** represent the same as in panels A and B for mice injected with aIgG2b liposomes.

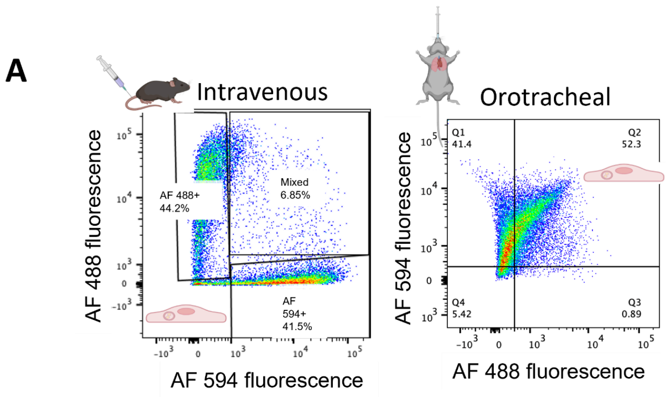


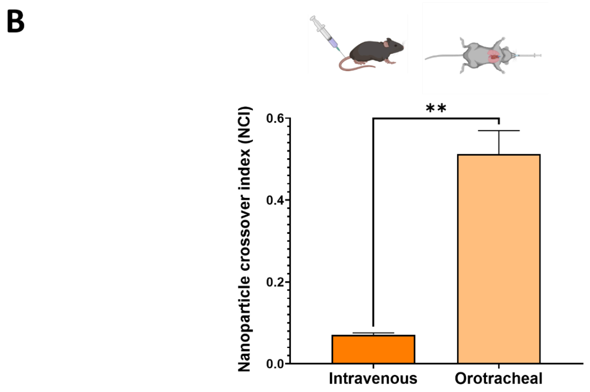


**Supp Fig 5: NCI positive control**. Comparison of intravenous injection and orotracheal instillation of liposomes. Orotracheal or intratracheal delivery of liposomes was used as a positive control. We have recently shown that when delivered in this way, liposomes remain in the airspace of the lungs and are not taken up by cells. This results in a high NCI ratio of 51.2% as shown in **A** (right panel) and **B.** Quantified using the NCI ratio

| Cell Type | Marker | Fluorophore | Dilution | Vendor |
|---|---|---|---|---|
| Leukocytes | CD45 | APC | 1:300 | Biolegend |
| Neutrophils | Ly6G | AF700 | 1:300 | Biolegend |
| Monocytes and Macrophages | CD64 | PE-Cy7 | 1:300 | Biolegend |
| Endothelial Cells | CD31 | PE-Cy7 | 1:250 | Biolegend |
| Epithelial Cells | EpCAM | BV711 | 1:250 | Biolegend |

| Cell Type | Markers used for identification |
|---|---|
| Neutrophils | CD45+, CD64-, Ly6G+ |
| Monocytes and Macrophages | CD45+, Ly6G-, CD64+ |
| Other Leukocytes | CD45+, Ly6G-, CD64- |
| Endothelial Cells | CD45-, EpCAM-, CD31+ |
| Epithelial Cells | CD45-, CD31+, EpCAM+ |
| Mesenchymal Cells | CD45-, CD31-, EpCAM- |

| Marker | Fluorophore | Dilution | Vendor |
|---|---|---|---|
| CD45 | FITC | 1:100 | Biolegend |
| CD15 | BV421 | 1:100 | Biolegend |
| CD206 | PE-Cy7 | 1:100 | Biolegend |
| CD31 | AF700 | 1:100 | Biolegend |
| EpCAM | PerCP/Cy5.5 | 1:100 | Biolegend |

| Cell Type | Markers used for identification |
|---|---|
| Neutrophils | CD45+, CD206-, CD15+ |
| Monocytes and Macrophages | CD45+, CD15-, CD206+ |
| Other Leukocytes | CD45+, CD15-, CD206- |
| Endothelial Cells | CD45-, EpCAM-, CD31+ |
| Epithelial Cells | CD45-, CD31+, EpCAM+ |
| Mesenchymal Cells | CD45-, CD31-, EpCAM- |

**Supp Table 3.** Fluorophores are used to identify cell types in murine models, as well as dilutions and vendors. **Supp Table 4**. Surface markers are used for the identification of murine cell types. **Supp Table 5**. Fluorophores and dilutions are used for the identification of human cells. **Supp Table 6**. Surface markers are used for the identification of cells obtained from *ex vivo* human lungs.

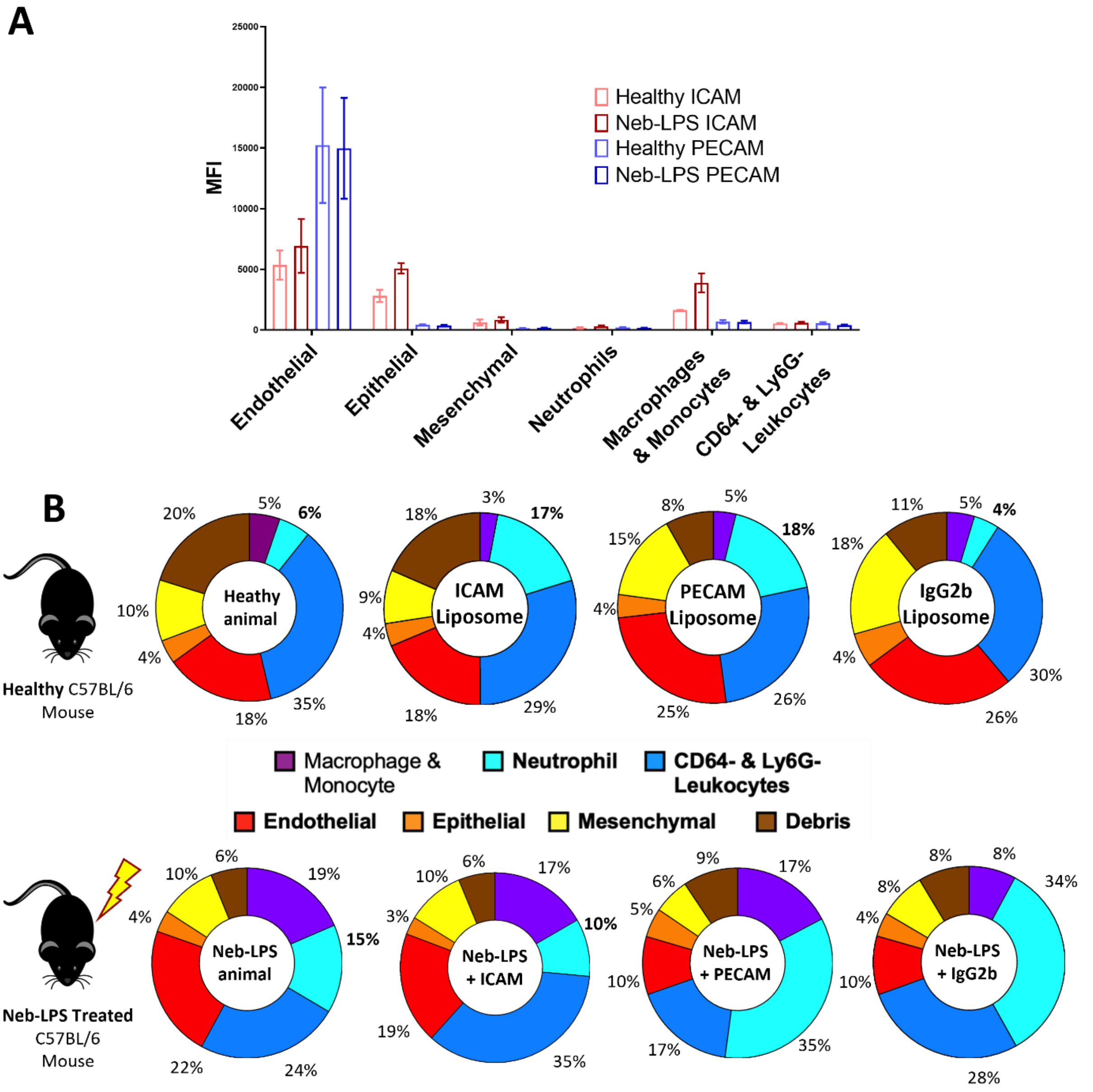


**Supp Fig 7. Changes in ICAM and PECAM receptors on different cell types in response to LPS injury and cell type distributions. A.** Expression of ICAM and PECAM on the surface of different cell types and changes in response to LPS stimulation. Overall, endothelial cells have the highest expression of these two cell markers. This does not change in response to LPS injury, with PECAM being more prominent on the endothelial surface. Interestingly, in response to injury, epithelial cells, macrophages, and monocytes show increased ICAM expression. **B.** Cell type distributions in a healthy and neb-LPS model of inflammation.

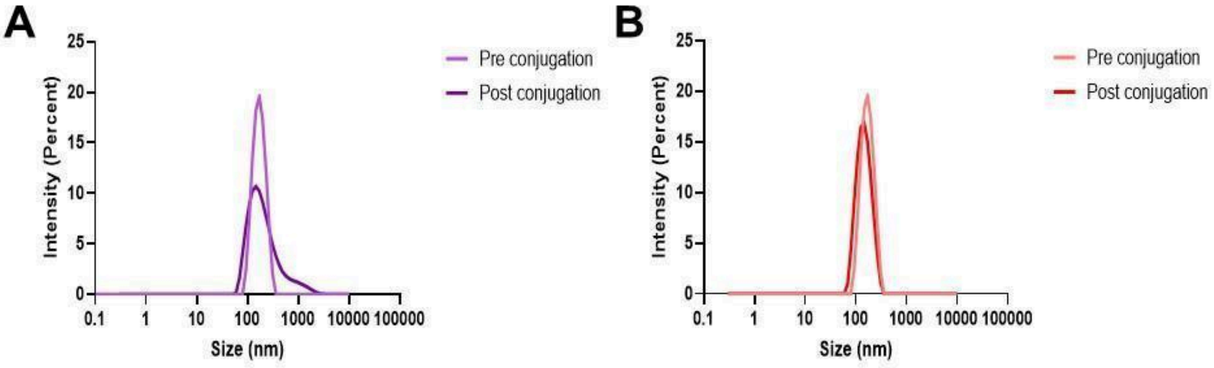


C

| Parameter | Pre-Conjugation | Post - Conjugation PECAM | Post-Conjugation IgG2 |
|---|---|---|---|
| Z average (nm) | 160.9 | 168.2 | 139.2 |
| Polydispersity index (PDI) | 0.075 | 0.2397 | 0.1338 |
| Peak mean by intensity | 174.5 | 274.1 | 150.9 |

**Supp Fig 8. Characterization of liposomes conjugated with PECAM and IgG2β monoclonal antibodies**. **A**. PECAM-conjugated liposomes demonstrate slightly more aggregation post-conjugation than IgG2b-conjugated liposomes (**Panel B**). **C**. Table representing the changes in nanoparticle sizes pre- and post-conjugation of the two different immunoliposomes.

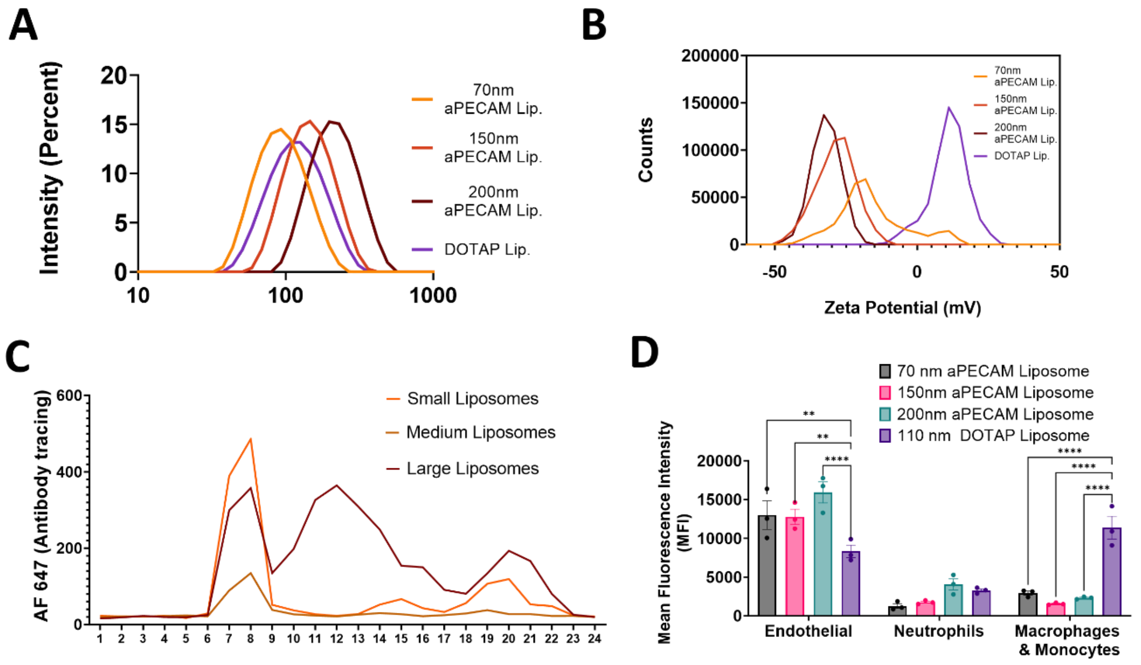


E

| Liposome type | Size (DLS) (nm) | PDI | Zeta Potential (mV) | Conjugation Efficiency |
|---|---|---|---|---|
| aPECAM small | 85.7 | .1417 | -16 | 90% |
| aPECAM med | 157.1 | .1317 | -26 | 80% |
| aPECAM large | 200.7 | .193 | -28 | 60% |
| DOTAP liposome | 108 | .136 | 10 | N/A |

F

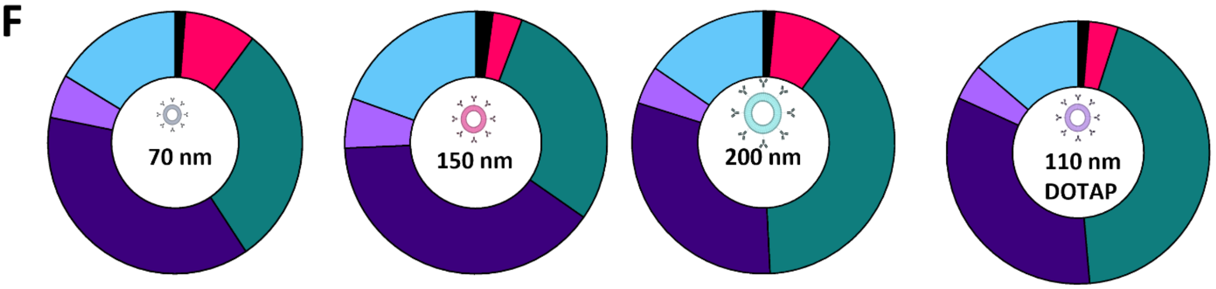


| Cell Type | 70nm | 150nm | 200nm | 100 nm DOTAP |
|---|---|---|---|---|
| Monocytes and Macrophages | 1.2 | 1.9 | 1.3 | 1.2 |
| Neutrophils | 8.4 | 3.3 | 8.0 | 3.3 |
| Leukocytes | 27.8 | 26.2 | 36.6 | 40.1 |
| Endothelial | 34.5 | 36.0 | 28.6 | 30.5 |
| Epithelial | 5.0 | 5.6 | 4.4 | 4.1 |
| Mesenchymal | 15.2 | 17.7 | 14.4 | 12.6 |

**Supp Fig 9: Characterization of varied size aPECAM liposomes and DOTAP liposomes. A.** Intensity plots depicting nanoparticle size distribution. **B.** Potential graphs showing the charge of differently sized and DOTAP liposomes. **C.** graph of nanoparticle conjugation efficiency fractions 6-9 are known to contain our liposomes, a peak here indicates correlation of antibody conjugation with nanoparticle. **D.** Mean fluorescence intensity (MFI) following uptake flow cytometry shows subtle changes in neutrophil uptake but high liposome fluorescence in monocytes and macrophages for DOTAP liposomes. **E.** Table describes the size, PDI, Zeta Potential, and conjugation efficiency of these liposomes. **F.** Cell type distribution does not significantly change with nanoparticle administration, regardless of size and charge.

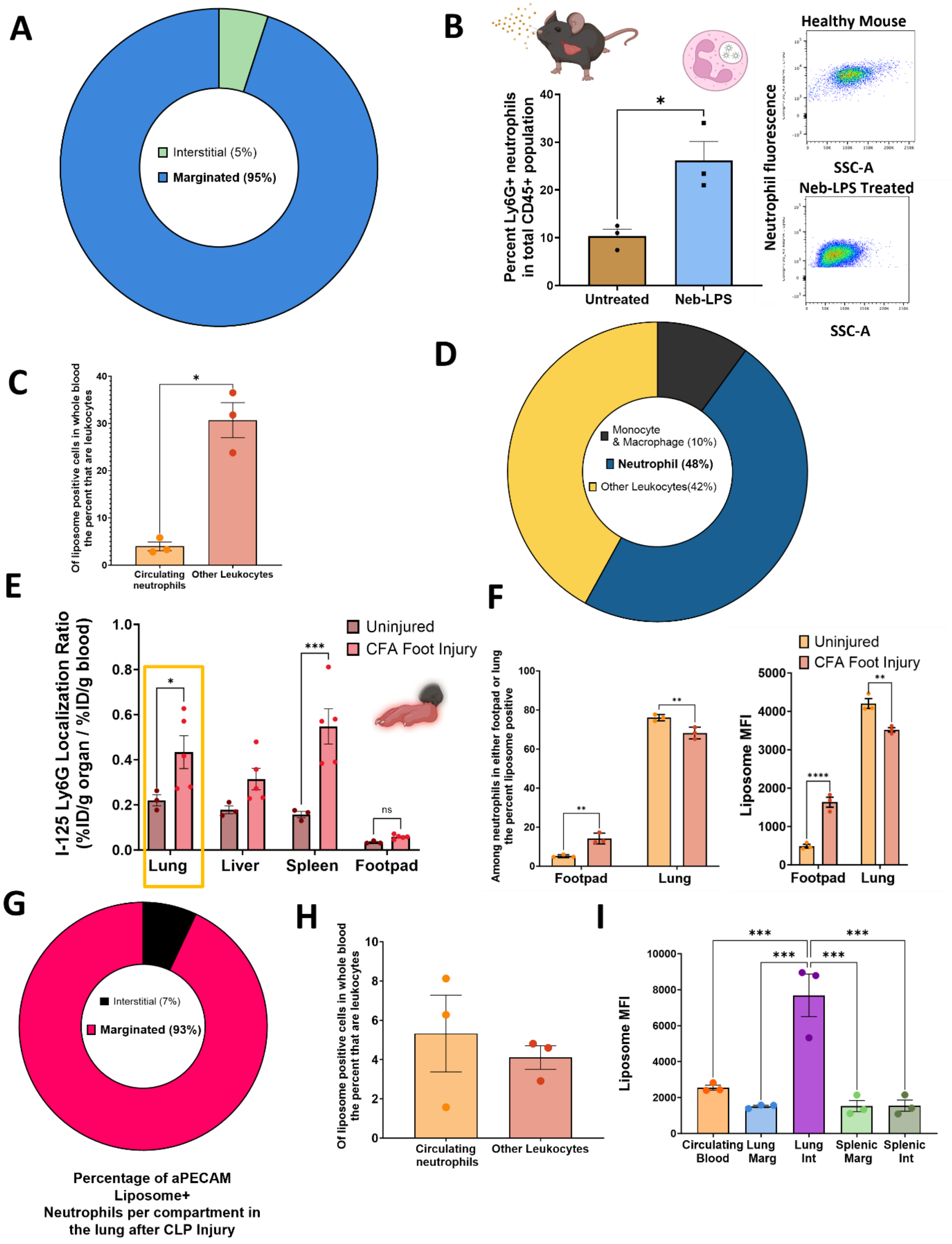
A
Interstitial (5%)
Marginated (95%)
B
Healthy Mouse
Neb-LPS Treated
SSC-A
Neutrophil fluorescence
Percent Ly6G+ neutrophils in total CD45+ population
Untreated
Neb-LPS
C
Of liposome positive cells in whole blood the percent that are leukocytes
Circulating neutrophils
Other Leukocytes
D
Monocyte & Macrophage (10%)
Neutrophil (48%)
Other Leukocytes(42%)
E
I-125 Ly6G Localization Ratio (%ID/g organ / %ID/g blood)
Uninjured
CFA Foot Injury
Lung
Liver
Spleen
Footpad
ns
F
Among neutrophils in either footpad or lung the percent liposome positive
Liposome MFI
Footpad
Lung
G
Interstitial (7%)
Marginated (93%)
Percentage of aPECAM Liposome+ Neutrophils per compartment in the lung after CLP Injury
H
Of liposome positive cells in whole blood the percent that are leukocytes
Circulating neutrophils
Other Leukocytes
I
Liposome MFI
Circulating Blood
Lung Marg
Lung Int
Splenic Marg
Splenic Int

**Supp Fig 10: Characterization of neutrophils in various models of inflammation A.** By flow cytometry, we first gate among total cells and then identify interstitial and marginated compartments. Gating this way shows that 95% of recovered neutrophils following disaggregation are marginated neutrophils. **B.** Neutrophils are highly present in the lung after nebulized LPS (neb-LPS) injury, as flow cytometry shows. **C.** Gating first by liposome-positive cells, we identify the liposome-positive circulating neutrophils fraction. **D.** Percentage breakdown of leukocytes after neb-LPS injury, shows that 48% of total leukocytes are marginated neutrophils. **E.** I-125 aLy6G biodistribution of neutrophil following injury by Complete Freund's Adjuvant (CFA) in the lung, liver, spleen, and affected paw show increased neutrophil margination in the lung and spleen. **F.** Flow cytometry and MFI of the footpad and lung showed that the lungs take up more aPECAM liposomes compared to the injured footpad and, as expected, take up more nanoparticles after injury, with a constitutive decrease in the lung. **G.** Following cecal ligation and puncture (CLP), we show that most liposome-positive neutrophils (93%) of liposome-positive neutrophils in the lung are in the marginated pool. **H.** Using the same gating strategy in blood, we find that about 5% of total liposome-positive cells are neutrophil-positive. **I.** Even though interstitial neutrophils represent a small fraction of total neutrophils in the lungs they take up a large portion of aPECAM liposomes compared to other neutrophil compartments assessed.

**A**

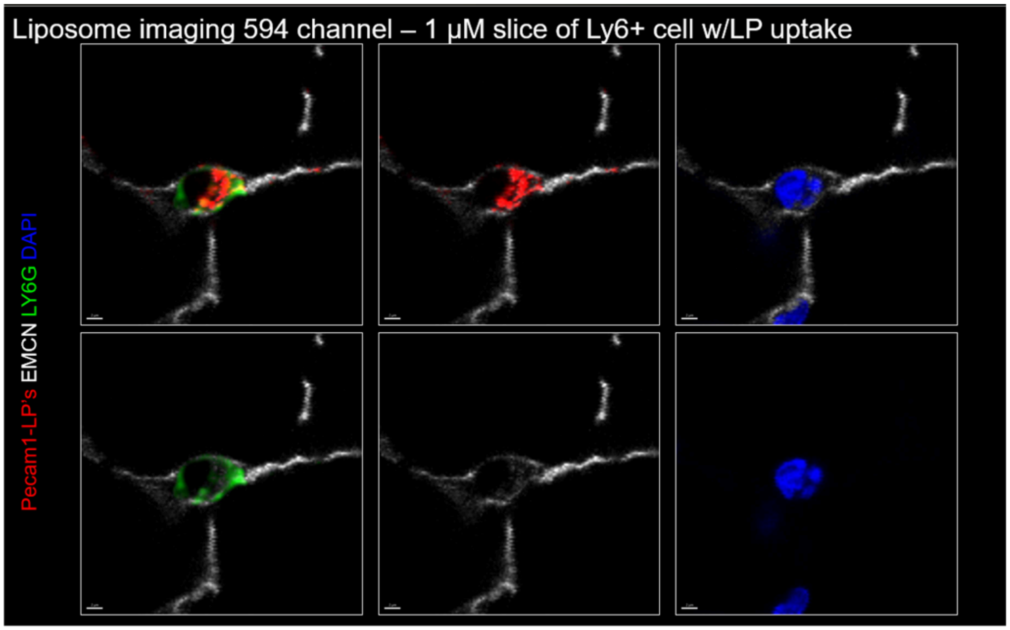


**B**

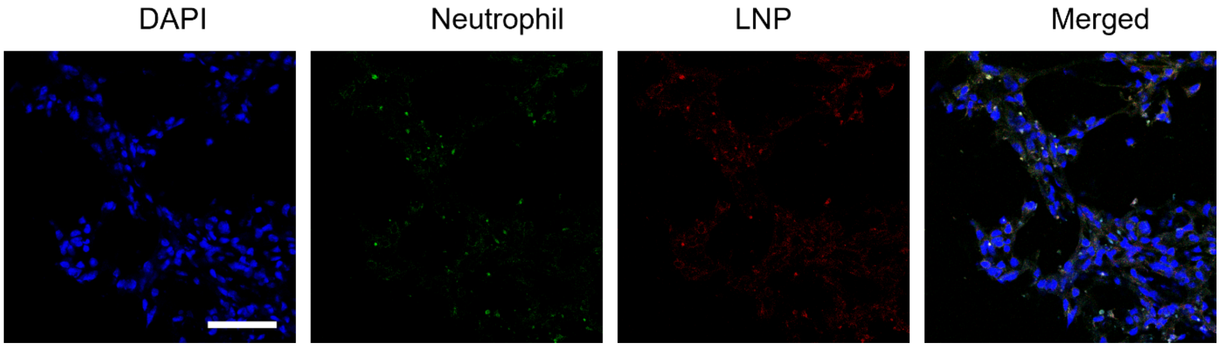


**Supp Fig 11: Microscopy single stains. A.** Single staining for thick-cut 3D histology samples. **B.** Single staining for confocal microscopy images of the lung after cecal ligation puncture (CLP) injury.

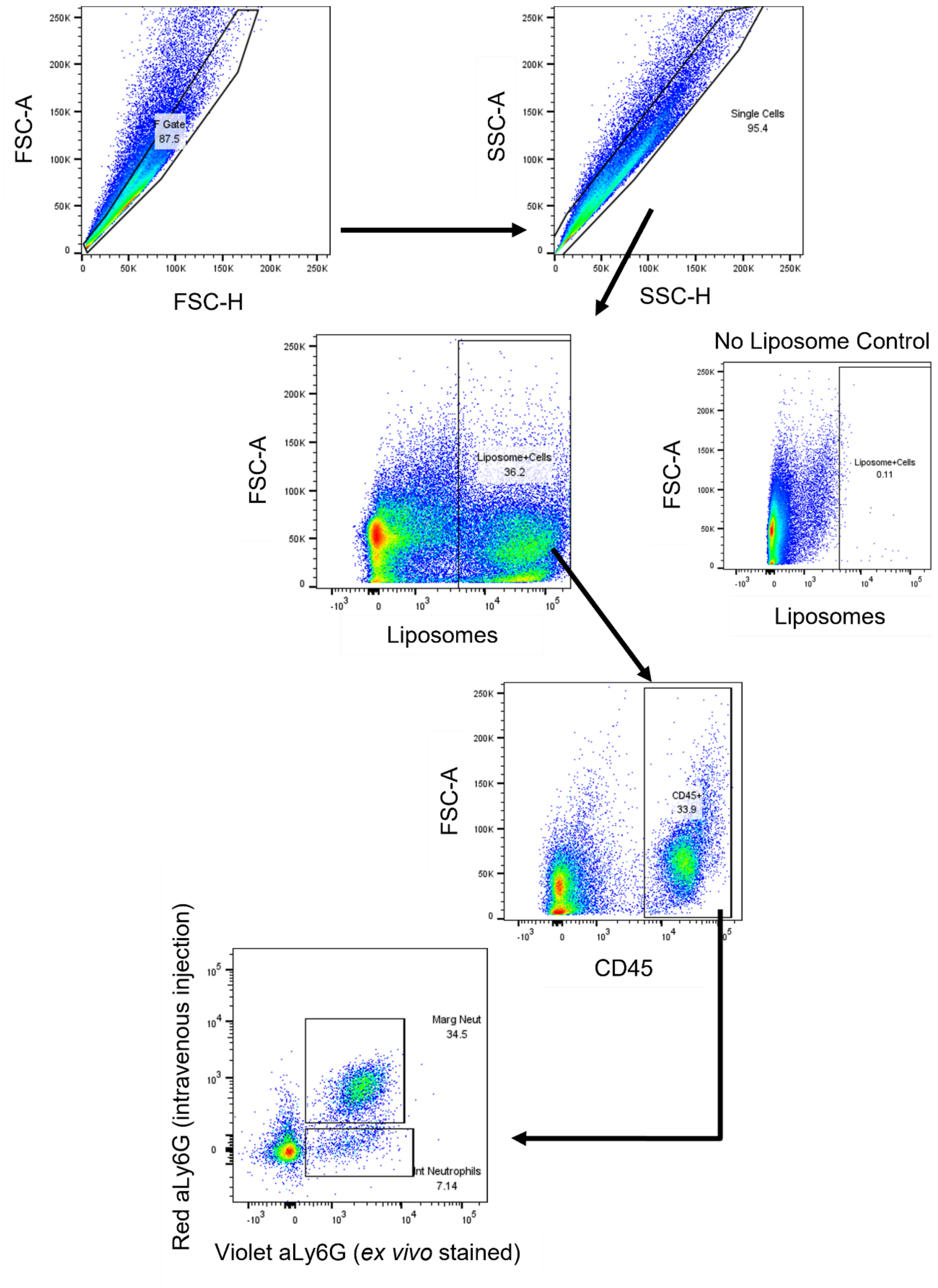


**Supplemental Fig 12:** Gating strategy used to determine marginated and interstitial neutrophils in the lung *in vivo.*

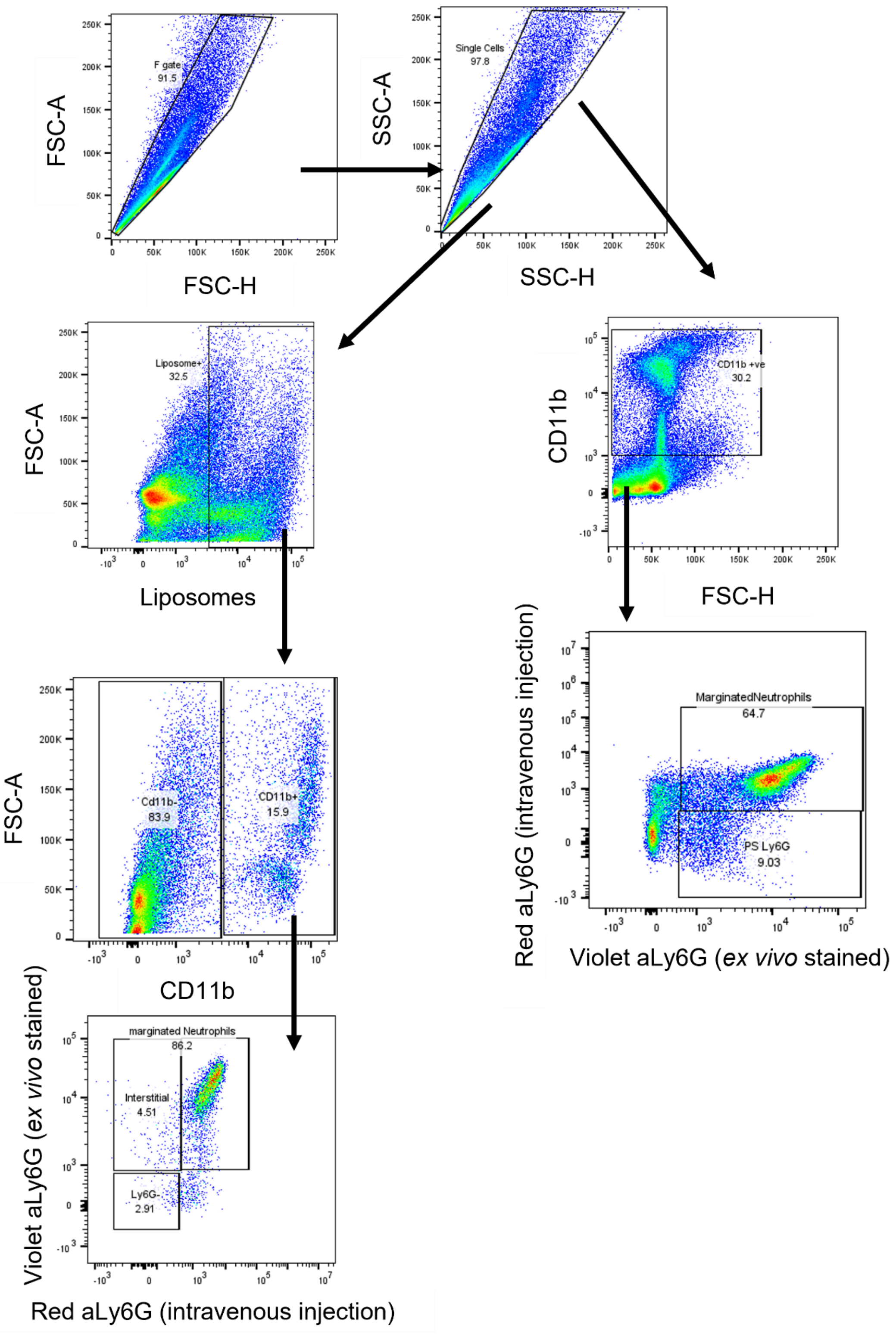


**Supplemental Figure 13:** Gating strategy used for gating lung, spleen and whole blood after cecal ligation and puncture (CLP)

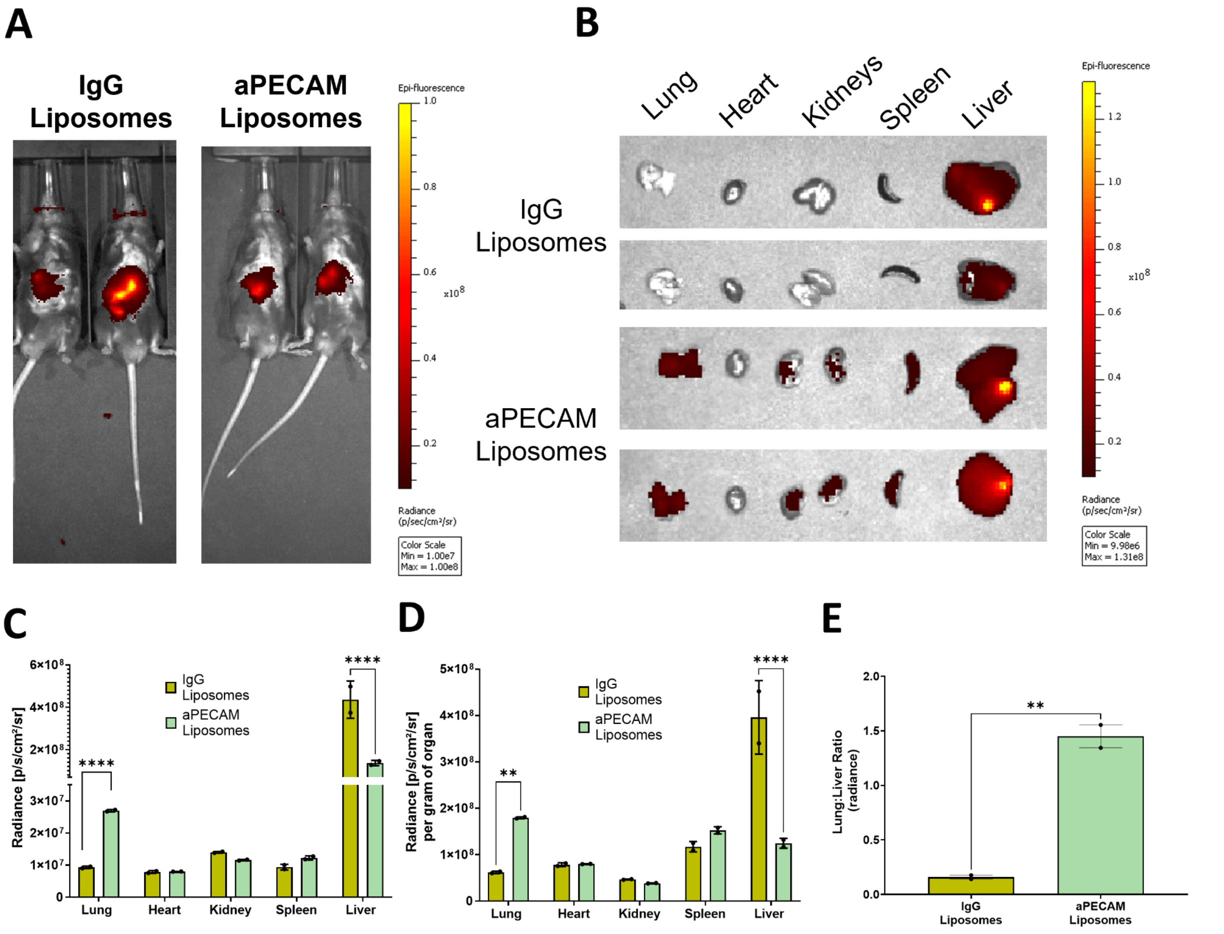


**Supplemental Figure 14: In vivo imaging system (IVIS) Biodistribution A.** *In vivo* whole animal imaging of mice injected with indocyanine green (ICG) loaded liposomes. Briefly, IgG and aPECAM liposomes were generated by thin film method; the film was hydrated in 500 uL of a 10 ug/mL ICG in 1xPBS. Liposomes were conjugated with either IgG or anti-PECAM antibody, and buffer was exchanged to remove free unloaded ICG. Mice were then injected at a dose of 10mg/kg by lipid mass and allowed to circulate for 30 minutes.**B.** *Ex vivo* imaging of organs taken immediately after *in vivo* imaging. aPECAM liposomes localize to the lung compared to IgG liposomes. **C.** Quantification of **B**, total radiance in each organ shows the preference of aPECAM liposomes to the lung compared to IgG, with a pairwise decrease in liver radiance. **D.** Total radiance normalized by organ weight to obtain the total radiance per gram of organ shows that aPECAM liposomes have higher uptake in the lung than in the liver relative to the mass of the organ. This is further reflected in the lung-to-liver ratio where a ratio higher than 1 shows preference to the lung over the liver.

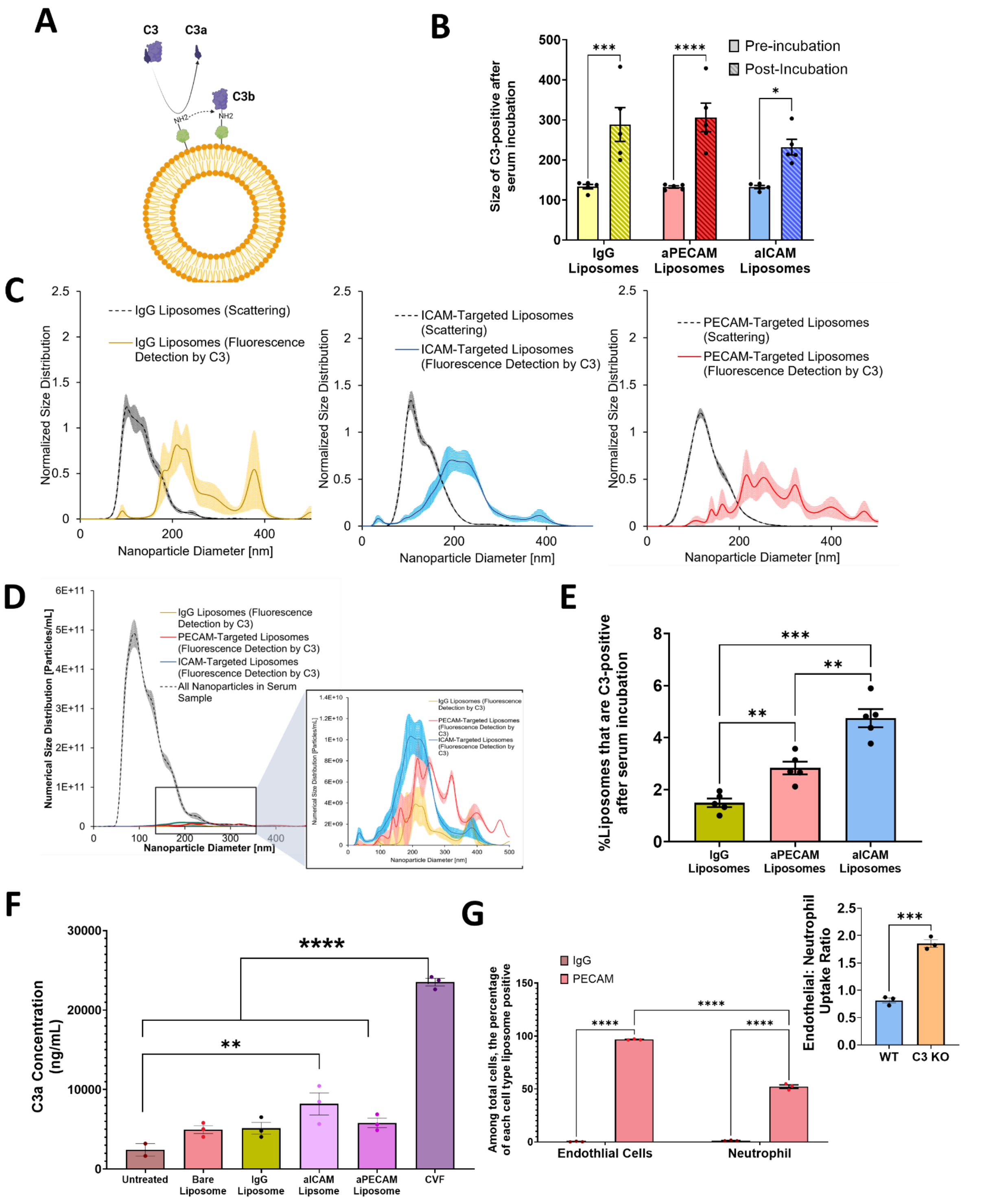


**Supplemental Figure 15: Complement deposition increases neutrophil recognition of aCAM-targeted nanocarriers. A.** Depiction of C3 deposition on nanoparticle surfaces in serum. Briefly, when C3 protein deposits on a foreign surface, it separates into C3a and C3b. C3b deposition on the surface is recognized by innate immune cells for clearance. C3a remains soluble in serum and can be used to catalyze further C3 activation; it is also implicated in complement-associated pseudoallergy (CARPA). **B.** Next, we assessed the effect of complement deposition on nanoparticle size. To test this, nanoparticle size was determined via nanoparticle tracking analysis (NTA) both before and after incubation in serum. C3b opsonized nanoparticles were assessed by spiking serum with a tracer concentration of fluorescently labeled C3 before incubation at 37 ℃. C3-bound particles were then recognized using a fluorescence filter that filters out any nanoparticle not fluorescently labeled by C3. **C**. Size distribution histograms normalized to particle number depict the change in size as shown in **B** for IgG, aPECAM, and aICAM liposomes. **D.** Size distribution of the IgG, aPECAM, and aICAM fractions targeted liposomes identified with fluorescent C3 protein on their surface compared to the size distribution of those, not C3 positive. **E.** This is quantified showing that while both aPECAM and aICAM-targeted liposomes had a higher percentage of C3-bound liposomes than IgG-conjugated liposomes, it still represents less than 10% of total incubated particles. **F.** We next tested serum incubated with either no liposomes, bare (no conjugation), IgG, aPECAM, or aICAM liposomes for the presence of C3a via ELISA assay. Bare, IgG, and aPECAM liposomes showed no significant increase in C3a concentration compared to control, aICAM liposomes show higher C3a and may be more inflammatory. **G.** We performed flow cytometry to understand the effect of complement deposition on neutrophil uptake of CAM-targeted nanoparticles. We injected aPECAM and IgG liposomes into C3KO mice and allowed circulation for 30 minutes. We observe no marginated lung neutrophils or endothelial cell uptake in mice treated with IgG liposomes. However, we observe an almost 50% reduction in the lung's neutrophil uptake of aPECAM liposomes. We then compare this to the endothelial-to-neutrophil uptake ratio of wild type (C57/B6) and show a 2.3-fold increase in endothelial preference in C3KO mice compared to wild type control.

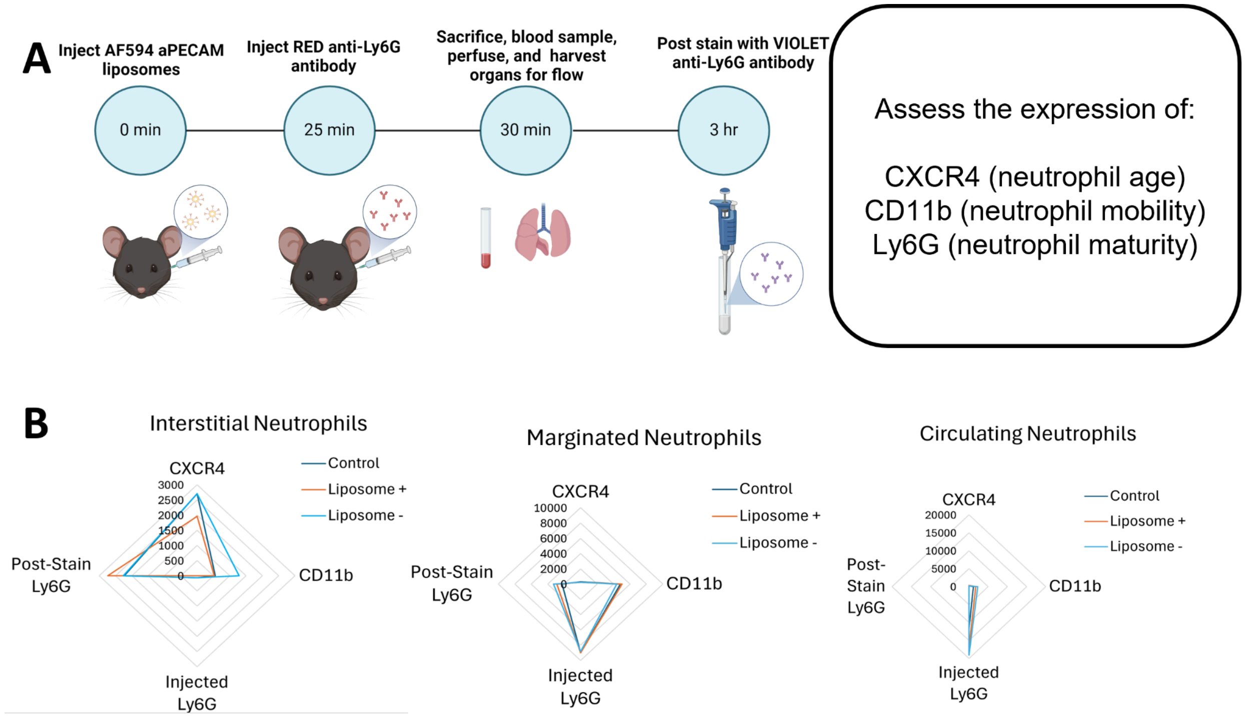


**Supplemental Figure 16: Surface marker presentation of neutrophil subtypes. A.** Schematic protocol for the identification of interstitial, marginated, and circulating blood neutrophils. After the isolation of neutrophil subpopulations, mice receiving aPECAM liposomes were further divided into liposome-positive and liposome-negative interstitial, marginated, or circulating neutrophils. These populations were then assessed for expression of CXR4, CD11b, and Ly6G via mean fluorescence intensity (MFI). **B.** Radar plots showing MFI of CXCR4, CD11b, and Ly6G in interstitial, marginated, and circulating neutrophils. We observe that interstitial neutrophils show elevated CXCR4 compared to both marginated and circulating neutrophils, while marginated neutrophils show increased CD11b compared to circulating neutrophils.

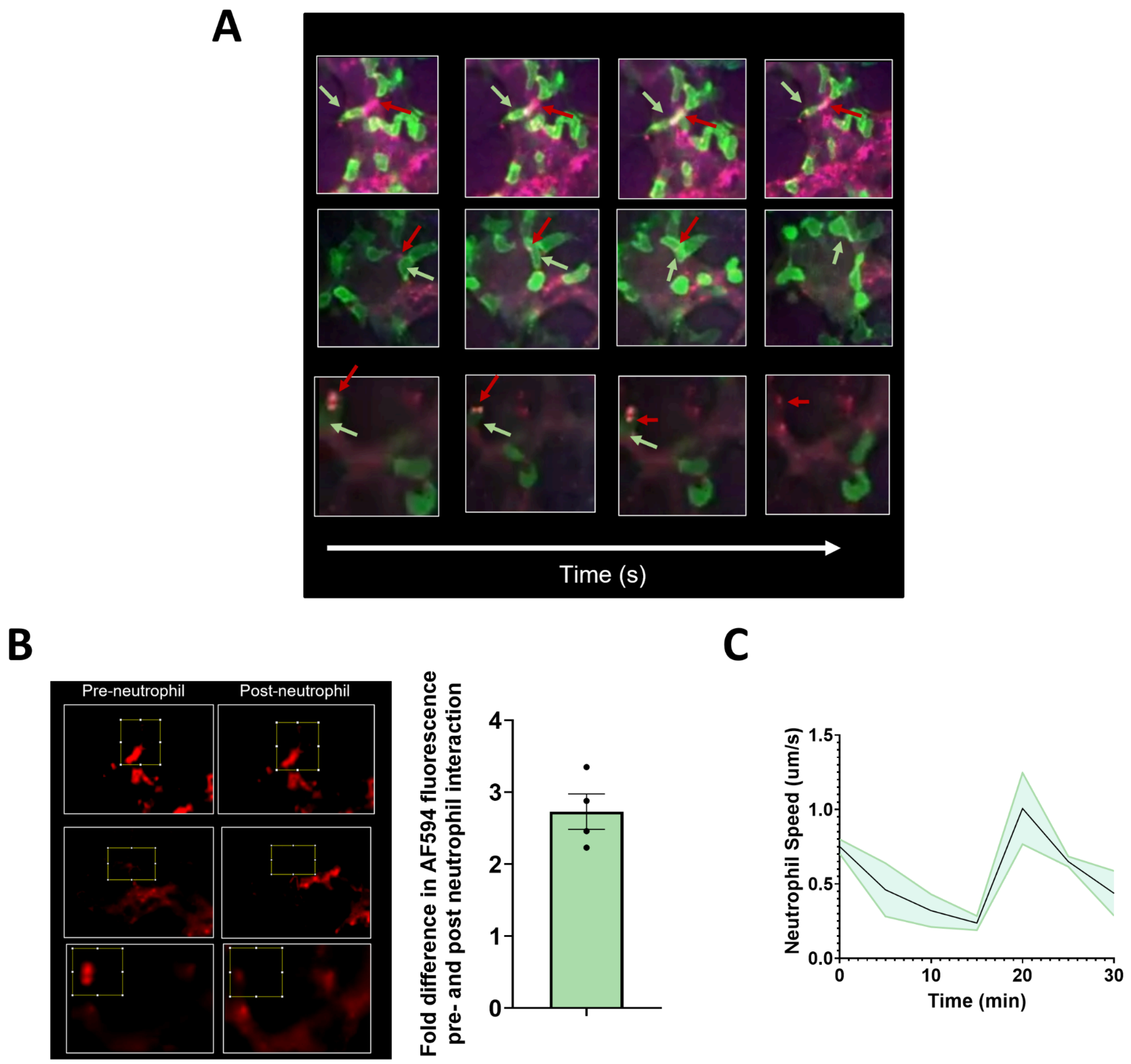


**Supplemental Figure 17** Additional analyses from videomicroscopy **A.** Additional stills demonstrating frame-by-frame uptake aPECAM liposomes by marginated neutrophils, stripping the liposomes away from endothelial cells. **B**. Region of interest (ROI) analysis of first frame versus final frame from **A**. Shows an almost 3-fold decrease in Red fluorescence within each ROI, showing neutrophil uptake and removal of liposomes from endothelial cells on capillary walls. **C.** Neutrophil crawl speed over time.

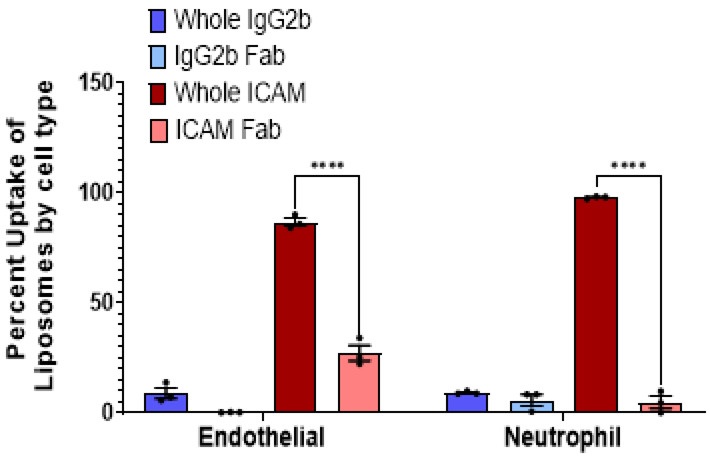


**Supplemental Figure 18:** Effects of removing fragment crystalizable region (Fc) on neutrophil and endothelial cell uptake of aICAM liposomes show a reduction in both but a near ablation of neutrophil uptake, suggesting Fc region recognition is a possible mechanism for neutrophil uptake of CAM-targeted nanoparticles.
